# Protein Aggregation Capture for Top-down Proteomics

**DOI:** 10.64898/2026.07.02.736076

**Authors:** Isabella G. Feltenstein, Bryon S. Drown

**Affiliations:** The James Tarpo Jr. and Margaret Tarpo Department of Chemistry, Purdue University, West Lafayette, IN, USA

**Keywords:** Proteoform, Top-Down Proteomics, Sample Clean-up, SP3

## Abstract

Proteins are dynamically regulated by a myriad of post-translational modifications (PTMs) that control their stability, conformation, activity, subcellular localization, and local interactions. Capturing the precise composition of these various modification states, or proteoforms, is a principal objective of top-down proteomics (TDP). By ionizing intact proteoforms and combining measurements of precursor ion and fragment ion masses, the position, stoichiometry, and combination of PTMs can be determined. Despite the highly valuable measurements that TDP can provide, it is typically less sensitive than corresponding peptide-level analysis with many reports utilizing input material in the microgram to milligram range.^1,2^ Contributing to this lack of sensitivity is the risk of sample loss due to non-specific binding to surfaces during sample preparation. The most widely employed sample preparation approaches for TDP either require high sample input (e.g. precipitation and ultra-filtration) or fail to effectively remove surfactants (e.g. solid-phase extraction). These limitations have hindered advancement of targeted TDP applications involving immunoprecipitation and other enrichment strategies. Bead-assisted protein aggregation, also referred to as single-pot, solid-phase-enhanced sample preparation (SP3), has emerged as a popular sample preparation strategy for bottom-up proteomic workflows,^3,4^ but has only been used in TDP with secondary ion exchange chromatography cleanup.^5^ We envisioned a magnetic bead based protein cleanup approach that proceeds directly to MS analysis with judicious choice of bead surface chemistry and elution conditions. Here we report a sample preparation method using hydroxyl-functionalized magnetic beads for top-down proteomics applications.

## Introduction

Top-down proteomics (TDP) is a powerful approach to identify proteoforms relevant to human health and disease that has advanced considerably with improved instrumentation. While discovery-oriented experiments have proven effective for broadly surveying the proteome, only a handful of proteoforms are typically confidently annotated per gene.^6–9^ Targeted TDP is a more focused variant in which proteoform families are enriched via immunoprecipitation against an endogenous epitope or genetically incorporated tag leading to significantly deeper characterization. This targeted approach has been successful in defining the proteoform landscape of KRAS,^10–12^ alpha-synuclein,^13^ cardiac troponin,^14^ apolipoproteins,^15,16^ and transthyretin.^17^ Mass spectrometry characterization of immunoglobulin repertoires has also emerged as a technique to understand the immune response of patients and sequence endogenous antibodies.^18–21^ Recent advancements in mass spectrometry instrumentation have enhanced these targeted strategies. In particular, Orbitrap-based charged detection mass spectrometry^22^ has extended the practical mass range of intact measurements and deconvolved overlapping fragment ions to enable precise measurement of larger and combinatorially modified proteoforms like MEK1,^23^ p53,^24^ tau,^25^ catenins,^26^ and endogenous antibodies.^27–30^ These studies represent an expansion in the capacity for targeted top-down mass spectrometry to make precise measurements and increasingly sophisticated data interpretation frameworks.

Despite these successes, TDP continues to face several significant limitations that stem from challenges in sample preparation. Matrix components such as surfactants and buffers that aid in analyte recovery pose problems at measurement time via ion suppression and chromatographic distortion. Depending on the type of interferent, the process of removal risks increased variability or sample loss.^1^ While surfactants play an important role in improving hydrophobic protein solubility, they are particularly critical in targeted TDP experiments. Non-ionic surfactants (e.g. Triton X-100 and NP-40) meaningfully reduce non-specific capture during the target enrichment step, and anionic surfactants (e.g. sodium dodecyl sulfate, SDS) provide suitable conditions for target elution. However, even trace amounts of SDS leftover in the final sample can suppress ionization during MS analysis, decreasing signal intensity and obscuring the presence of low-abundance proteoforms.^31^ This incompatibility creates an inherent trade-off: while surfactants enable efficient solubilization and specific enrichment, their removal is essential before MS analysis.

A variety of sample preparation approaches for TDP have been introduced by the community, each with their own merits and limitations. Protein precipitation with an organic solvent or at an aqueous-organic biphasic interface (e.g. methanol-chloroform-water, MCW) is a venerable approach for TDP that is effective at removing ionic detergents.^32–35^ While some prior studies have shown that it is biased towards the recovery of larger proteins and proteoforms,^36^ it is widely employed in discovery proteomics applications due to its compatibility with gel-based fractionation methods.^37,38^ Although broadly used, this approach is technically challenging to execute with limited sample input and is not amenable to automation. Other approaches including size exclusion chromatography,^39^ ultrafiltration,^40^ and solid-phase extraction^41^ can be effective but struggle to remove surfactant micelles. Alternatively, degradable surfactants or species with limited ion suppression can obviate the need for their removal, however these species do not offer the same denaturing capacity as SDS.^42–44^ Bead protein aggregation capture, often referred to as SP3, has proven to be a versatile approach for bottom-up proteomics studies due to its broad tolerance for interferents and ability to concentrate trace inputs.^45,46^ Adaptation of SP3 for TDP have utilized carboxylate-functionalized bead surfaces which have necessitated protein resolubilization in concentrated formic acid or dilute SDS.^5,47^ Consequently, a gap remains for single-step protein sample preparation that can tolerate the interferents present in targeted TDP experiments.

Given the potential promise of SP3-based workflows to significantly concentrate samples while removing a wide range of interferents, we sought to develop a sample preparation method for TDP that involved protein aggregation on magnetic beads for direct recovery for RPLC-ESI-MS. In initial development of this method, we evaluated different bead surface chemistries and recovery buffers on standard protein mixes and cell lysate. In the process, we investigated the extent of SDS removal using the ContamSpot assay and performance with highly dilute samples. Overall, we demonstrate the suitability of a single-step approach for protein purification using bead aggregation capture and acidic elution condition.

## Experimental Section

### Materials

Reagents, protein standards, magnetic beads, and consumables with suppliers and catalog numbers are listed in Table S1. The Pierce Intact Protein Standard Mix (Thermo Fisher), and a four-protein mix of myoglobin, carbonic anhydrase, trypsinogen, and ubiquitin (NRTDP standard) as defined proteoform mixtures during method optimization.

### Magnetic Bead Sample Preparation

Three clean-up strategies were compared: methanol-chloroform-water (MCW) precipitation, SP3 with carboxylate-functionalized beads (Cytiva), and SP3 with hydroxyl-functionalized beads (MagReSyn). In the hydroxyl-bead method, protein samples were reduced with DTT (10 mM, 60 °C, 30 min), combined with hydroxyl beads (100 µg) and acetonitrile to 70% (v/v), and aggregated by mixing at 1000 rpm for 10 min at room temperature. Beads were collected on a magnetic stand, washed twice with 100% acetonitrile without disturbing the bead-protein pellet, and proteins were eluted directly into MS-compatible solvent with 0.5% TFA (1000 rpm, 20 min). For comparison, the Cytiva method used 50% ethanol for binding, 80% ethanol washes, and 0.5% TFA elution; MCW precipitation followed a standard protocol^33^ with final resuspension in 5% acetonitrile/0.2% formic acid. Full step-by-step protocols for all three methods including standard protein mix (4.5 µg) and lysate inputs are provided in the Supporting Information (Section S2).

### Cell Culture and Lysate Preparation

HeLa cells were cultured in supplemented DMEM, harvested by scraping at passage 15, and lysed by probe sonication in 50 mM HEPES (pH 8.5), 1% SDS, 1× Halt protease inhibitor. Clarified lysate (∼2 mg/mL) was reduced with DTT (10 mM, 60 °C, 30 min) and aliquoted (30 µL, ∼60 µg per condition) for clean-up by each of the three methods. Detailed procedures are given in the Supporting Information (Sections S3–S4).

### LC-MS/MS Analysis

Purified proteins were analyzed by reversed-phase LC-MS/MS on a Vanquish Neo hyphenated to an Orbitrap Exploris 480 (Thermo Fisher) operated in intact-protein mode. Proteins were separated on a MAbPac capillary RP column (150 µm i.d. x 15 cm L) at 1.5 µL/min with buffer A: 0.1% formic acid in water; buffer B: 0.1% formic acid in acetonitrile. Standard and lysate samples used a 4–64% B gradient over ∼55 min. MS1 spectra were acquired at 120,000 resolving power with a scan range 500–2000 *m/z*, and data-dependent HCD MS2 spectra (top-3) were acquired at 60,000 resolving power. Full acquisition parameters for mass spectrometry acquisitions are reported in Table S3.

### Detergent Removal Assay

Removal of SDS was assessed colorimetrically with the ContamSpot^48^ assay using BSA reconstituted in 50 mM HEPES, 1% SDS, before clean-up and after elution with either 100 mM ammonium bicarbonate or 0.5% TFA (v/v); a TFA-only reference spot controlled for ion-pair color formation. The full protocol is provided in the Supporting Information (Section S5).

### Dilute Sample Recovery

Equine myoglobin was dissolved and diluted to 4 ng/µL in 0.5% SDS (v/v), 50 mM HEPES (pH 7.4) and divided into 50 µL aliquots (200 ng). Each aliquot was subjected to either MCW or SP3-based cleanup. Protein samples were resuspended/recovered in 25 µL 2% acetonitrile, 0.1% formic acid, 97.9% water (MCW method) or 25 µL 0.5% TFA (SP3 method). Samples (5 µL injection) were analyzed with a MS1-only acquisition method.

### Data Analysis

Annotated proteoform and subsequence searches were performed with Proteome Discoverer 3.0 SP1 using the ProSight PD node (version 4.2)^49^ against the human SwissProt database (obtained 6/13/2023) with a high-resolution MS1/high-resolution MS2 (“high–high”) crawler, 2.2 Da precursor and 10 ppm fragment ion mass tolerances, and 1% FDR control at the spectrum, proteoform, and protein levels. Search results are provided as a tdReport file that is browsable with TDViewer (https://tdviewer.kelleher.northwestern.edu/). GRAVY scores and isoelectric points were computed from identified sequences using custom Python code (see Data Availability). For the dilute sample recovery experiment, myoglobin abundance was calculated with ProteoQuant (version 0.2.1) which provides apex intensity and extracted-ion chromatogram area-under-the-curve abundances for proteoform targets. Statistical significance of myoglobin intensity was performed by unpaired two-sided Welch’s *t*-test. Differences among physiochemical properties of proteoforms identified following each cleanup methods (grouped both by bead type and by individual condition) were evaluated for each property using Kruskal-Wallis tests with eta-squared (η²) effect sizes, followed by pairwise Mann-Whitney U tests with Cliff’s δ effect sizes and two-sample Kolmogorov-Smirnov tests. Pairwise p-values were corrected by the Benjamini–Hochberg procedure.

## Results and Discussion

### Evaluation of SP3-based Clean-up with Standard Protein Mixtures

Recovery of intact proteins using carboxylate-functionalized beads presents significant challenges, often requiring harsh elution conditions such as 80% formic acid.^47^ While solvent levels of acid are efficient at recovering proteins off these beads, proteins exhibit limited stability under these conditions. Seeking less harsh protein recovery conditions, we evaluated the ability of several commercially available magnetic beads to capture and prepare the Pierce Intact Protein Standard Mixture (Pierce) and the NRTDP Standard Mixture for mass spectrometry analysis (Figure 1, S1, S2, S3). Variable protein recovery was observed with each standard mix, depending on bead surface chemistry used for aggregation. The optimized conditions utilized hydroxyl-functionalized beads with 0.5% TFA recovery buffer (Figure 1C). All proteins were detected under these conditions with minimal alteration to their elution profiles. The signal-to-noise (S/N) ratio for the [M+23H]^23+^ precursor ion of Protein G was highest with methanol-chloroform-water (MCW) precipitation at 292.91 (Figure 1B) but was comparable using MagReSyn hydroxyl beads with 0.5% TFA as the recovery buffer at 212.96 (Figure 1C). In contrast, when using Cytiva carboxylate-functionalized beads with 0.5% TFA recovery buffer, the same precursor ion of Protein G exhibited a significantly lower S/N ratio of 57.51 (Figure 1D). These preliminary findings suggest that appropriate choice of magnetic bead surface chemistry can enable intact protein capture and recovery for mass spectrometry analysis.

**Figure 1.**
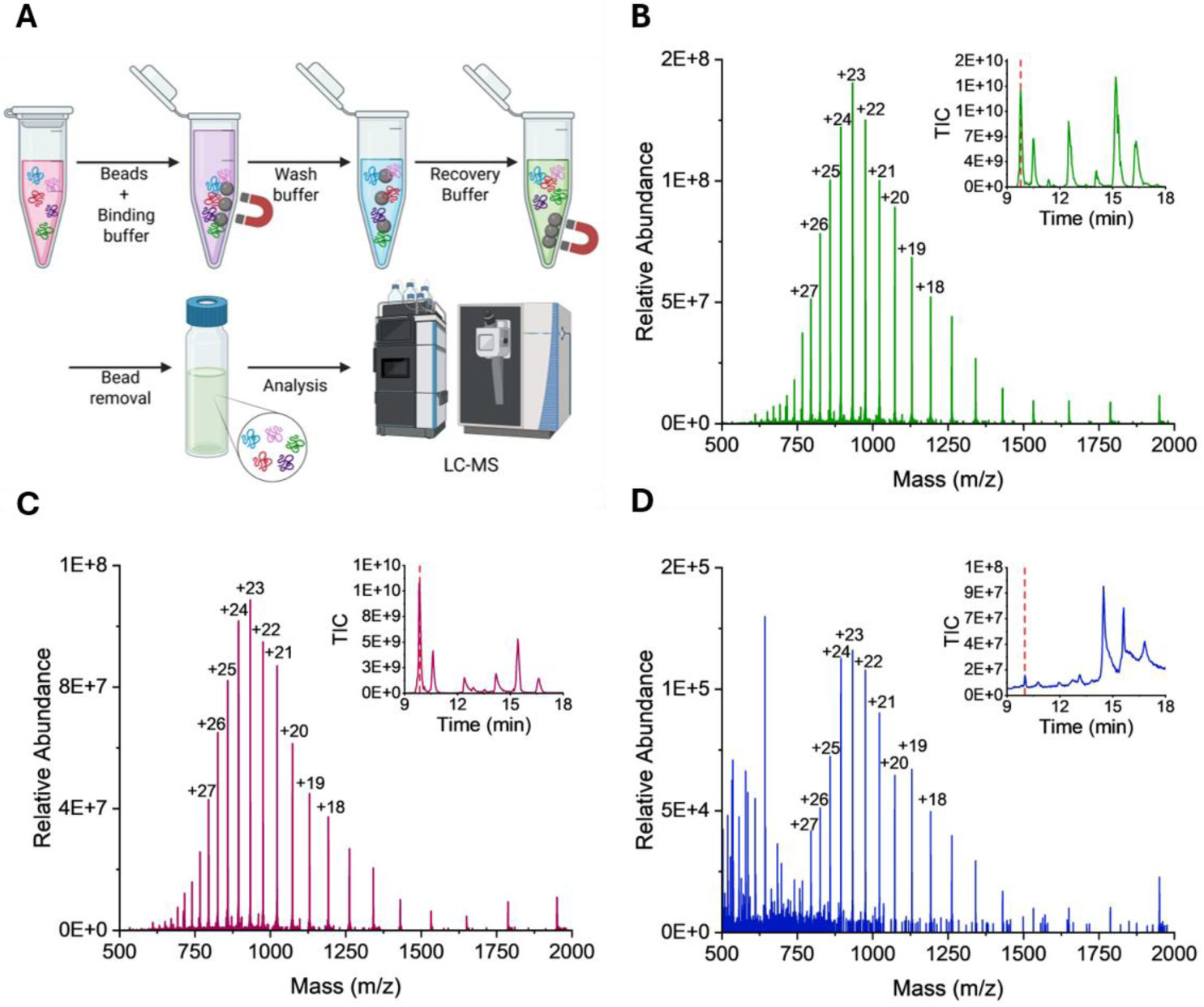
Evaluation of SP3-based cleanup of Pierce. (A) Schematic of bead aggregation workflow. Protein sample in buffer is combined with magnetic beads and binding buffer to facilitate protein aggregation on beads. Post-aggregation, the protein-bead complex is washed and then disrupted to recover proteins using a recovery buffer and analyzed using RPLC-MS. Representative intact single scan mass spectra of Protein G from each sample preparation technique. (B) Methanol-chloroform-water precipitation, (C) hydroxyl MagReSyn beads, and (D) carboxylate-functionalized Cytiva beads.

Encouraged by these results, we evaluated two critical factors in the utility of cleanup methods – sensitivity and ability to eliminate ionic detergents. To assess sensitivity, we carried out an experiment seeking to recover myoglobin from a dilute solution (4 ng/uL, 200 ng total) in the presence of 0.5% SDS by MCW or hydroxyl SP3. At this concentration, MCW precipitation produces a protein pellet that is not visible, making execution of this procedure technically challenging. The recovered protein was analyzed by LC-MS and quantified with ProteoQuant, an open-source application that fits isotopic distributions of target proteoforms to experimental data and returns apex and LC peak-integrated intensities (Figure 2A). While hydroxyl SP3 provided a 2.4- and 3.1-fold improvement over MCW in AUC and apex abundance, respectively, these improved recoveries were not significant. Nonetheless, these data suggest that the ease of hydroxyl SP3 sample preparation does not come at the expense of sensitivity.

**Figure 2.**
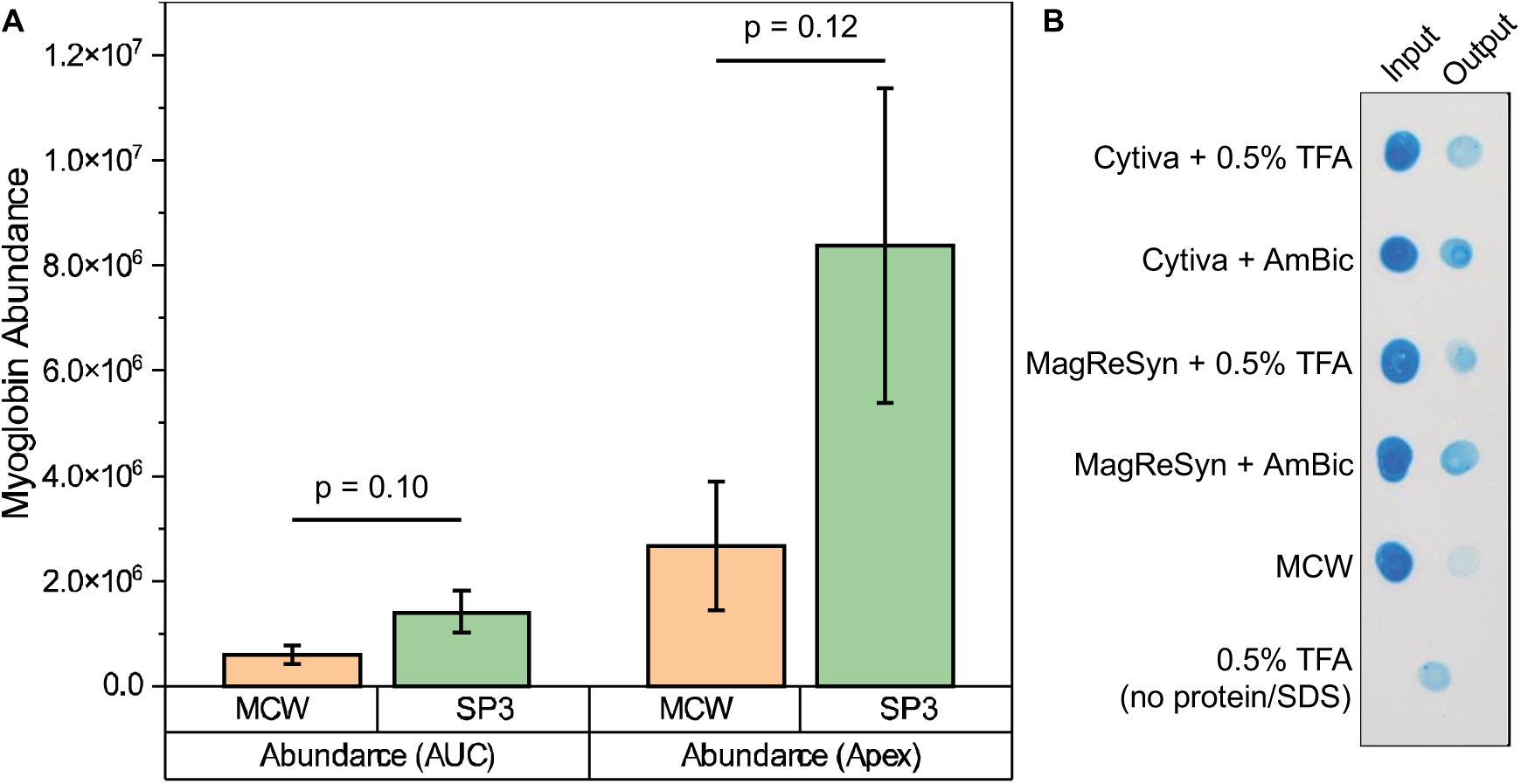
(A) Quantification of recovered myoglobin from dilute solution (4 ng/μL) via methanol-chloroform-water precipitation (MCW) or MagReSyn hydroxyl bead capture (SP3). Bar plot represents the mean and error bars represent standard error of the mean of six independent replicates. Reported p-values correspond to unpaired two-tailed Welch’s t-test. (B) ContamSpot assay results detecting presence of SDS following cleanup of BSA reconstituted in 1% SDS, 50 mM HEPES by various methods. Brighter blue dots represent higher concentration of ion pairing agent (SDS or TFA).

The effective removal of detergents from protein samples was measured using a ContamSpot colorimetric assay before and after clean-up.^48^ Starting with samples of bovine serum albumin (BSA) reconstituted in 50 mM HEPES and 1% SDS, we performed our clean-up procedure and evaluated the difference in SDS presence prior to clean-up and after elution with either ammonium bicarbonate or 0.5% TFA. In this assay, *O*-toluidine blue is added to the aqueous sample, washed with ethyl acetate, and the organic layer spotted on a TLC plate. Detergents and some ion-pairing agents act as phase-transfer catalysts leading to partitioning of the dye and enhanced intensity of color (Figure 2B). Comparison of pre- and post-cleanup shows removal of SDS following bead-based clean-up. Precise determination of remnant SDS is complicated by TFA strongly ion pairing with *O*-toluidine leading to similar color intensity from post-hydroxyl SP3 cleanup and 0.5% TFA alone. These preliminary findings with protein standards demonstrating sensitive recovery and removal of interferents provided a strong rationale for applying the bead-assisted aggregation method to a more complex and biologically relevant sample.

### Development of Cell Lysate Clean-up for TDP

While protein standards are helpful in establishing initial workflow conditions, the translational potential of this approach depends on its robustness with complex samples. To this end, HeLa cell lysate was prepared, and proteins were extracted with an SDS-based lysis buffer. This complex protein sample was subjected to the same three cleanup approaches described in the evaluation of protein standard mixtures (MCW, SP3 with MagReSyn hydroxyl beads, and SP3 with Cytiva carboxylate beads), and the resulting proteins were analyzed by single-shot RPLC-MS/MS using a high-high top-3 data dependent acquisition on an Exploris 480 (Figure 3A). In addition to varying bead surface chemistry, acidic recovery buffer identity and strength was varied. Annotated proteoform and subsequence searches were performed with ProSight PD and counts of protein and proteoform identifications are outlined in Table 1. In agreement with experiments performed with protein standards, these data suggest MagReSyn hydroxyl beads with 0.5% TFA was most performant with the highest number of proteoforms among bead cleanup methods and protein identifications overall (231 vs 228 for MCW). Proteoform identifications were distributed across retention times but were concentrated in the first half of the gradient, consistent with a correlation in hydrophobicity and proteoform size (Figure S4). SDS-PAGE analysis of boiled beads before and after acidic elution indicated that a limiting factor for this approach remains incomplete resolubilization of captured protein aggregates on the bead surface (Figure S5).

**Figure 3.**
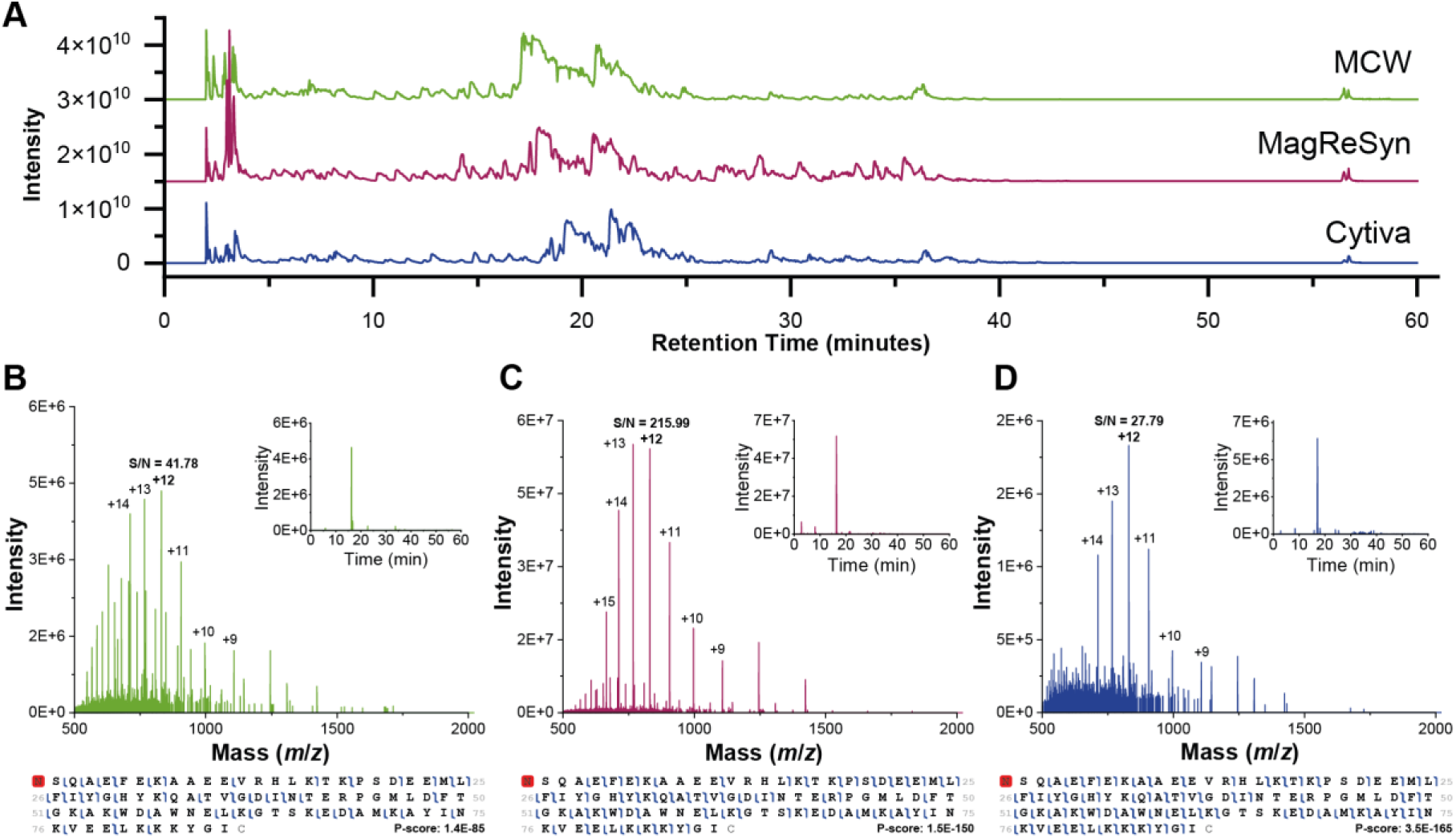
Comparison of cleanup approaches on HeLa cell lysate. (A) Stacked total ion chromatograms following each cleanup method. MS1 spectra with extracted ion chromatograms and fragment maps of primary proteoform of Acyl CoA binding protein across tested conditions, MCW (A), MagReSyn with 0.5% TFA (B), and Cytiva w/ 0.5% TFA (C).

**Table 1.** Proteins and proteoforms identified by top-down mass spectrometry.

| **Cleanup Method** | **Recovery Buffer** | **Proteins** | **Unique Proteoforms** | **Annotated Proteoforms** | **Subsequence Proteoforms** |
| --- | --- | --- | --- | --- | --- |
| MCW | 5% ACN, 0.1% FA | 228 | 735 | 480 | 429 |
| Cytiva | 10% FA | 185 | 544 | 372 | 291 |
| Cytiva | 20% FA | 158 | 506 | 337 | 264 |
| Cytiva | 0.5% TFA | 204 | 560 | 380 | 311 |
| Cytiva | 2% TFA | 105 | 431 | 276 | 262 |
| MagReSyn Hydroxyl | 10% FA | 206 | 572 | 405 | 306 |
| MagReSyn Hydroxyl | 20% FA | 186 | 470 | 325 | 259 |
| MagReSyn Hydroxyl | 0.5% TFA | 231 | 580 | 421 | 314 |
| MagReSyn Hydroxyl | 2% TFA | 173 | 486 | 321 | 298 |
| Total unique |  | 347 | 1385 | 843 | 847 |

Beyond identification counts, we examined whether the quality of proteoform characterization data varied with sample preparation methods and found that P-score distributions of shared proteoforms were broadly similar (Figure S6A, B). As precursor mass deconvolution is a key determinant in proteoform identification, we also examined the ability of the open source tool FLASHDeconv^50^ to perform deconvolution and mass inference, finding that MCW and hydroxyl MagReSyn with 0.5% TFA led to the highest enrichment of high Qscores (Figure S6C). Using a recently reported approach for estimating FDR in mass deconvolution,^51^ we observed these two cleanup methods yielded the greatest number of deconvoluted masses at 5% FDR (Figure S6D). This trend is exemplified by the most confidently identified proteoform in the dataset from Acyl-CoA-binding protein (PFR00000001054). We compared both the extracted ion chromatograms and inspected apex MS1 scans of this proteoform across conditions. Noticeable noise reduction and improved resolution of the hydroxyl-bead sample is evident compared to the carboxylate bead sample and MCW, with S/N ratios of the [M+12H]^12+^ species reported as 41.78 for MCW, 215.99 for MagReSyn, and 27.79 for Cytiva (Figure 3B-D). Together, these findings suggest that sample preparation with hydroxyl MagReSyn beads offers a comparable degree of proteoform characterization as MCW.

Given the different chemical environments in which proteins aggregate in each cleanup approach, we sought to determine the presence of any bias in the physiochemical properties of proteins recovered as others have observed. Initial evaluation of proteoforms by their GRAVY score, isoelectric point, and average mass was skewed due to extensive representation by histone proteoforms in the data set, specifically a seemingly biased result towards more basic proteoforms (Figure 4A). Follow-up analysis of non-histone proteoforms led to the emergence of a bimodal distribution in isoelectric point that typifies most proteomes (Figure S7). Both omnibus and pairwise hypothesis testing identified some significant differences in distributions, but the corresponding effect sizes were uniformly negligible to small. Grouped by bead type, no property exceeded an omnibus effect size of η^2^ = 0.008 (Kruskal-Wallis), and the same held when conditions were considered individually (η^2^ ≤ 0.018). In pairwise comparisons, every difference that remained significant after multiple-testing correction was small in magnitude (Cliff’s δ ≤ 0.25). Therefore, these results indicate that PAC-based cleanup recovers proteoforms spanning the same ranges of mass, charge, and hydrophobicity as the reference MCW workflow. To further investigate whether cleanup approaches favor particular chemical modifications, mass shift analysis was performed for the top conditions for each bead type compared to MCW (Figure 4B). Several common PTMs were observed such as methylation, oxidation, acetylation, and phosphorylation. All three of these primary methods avoided introducing artifactual modifications, however higher concentrations of organic acid led to the more frequent occurrence of unannotated mass shifts (Figure S8).

**Figure 4:**
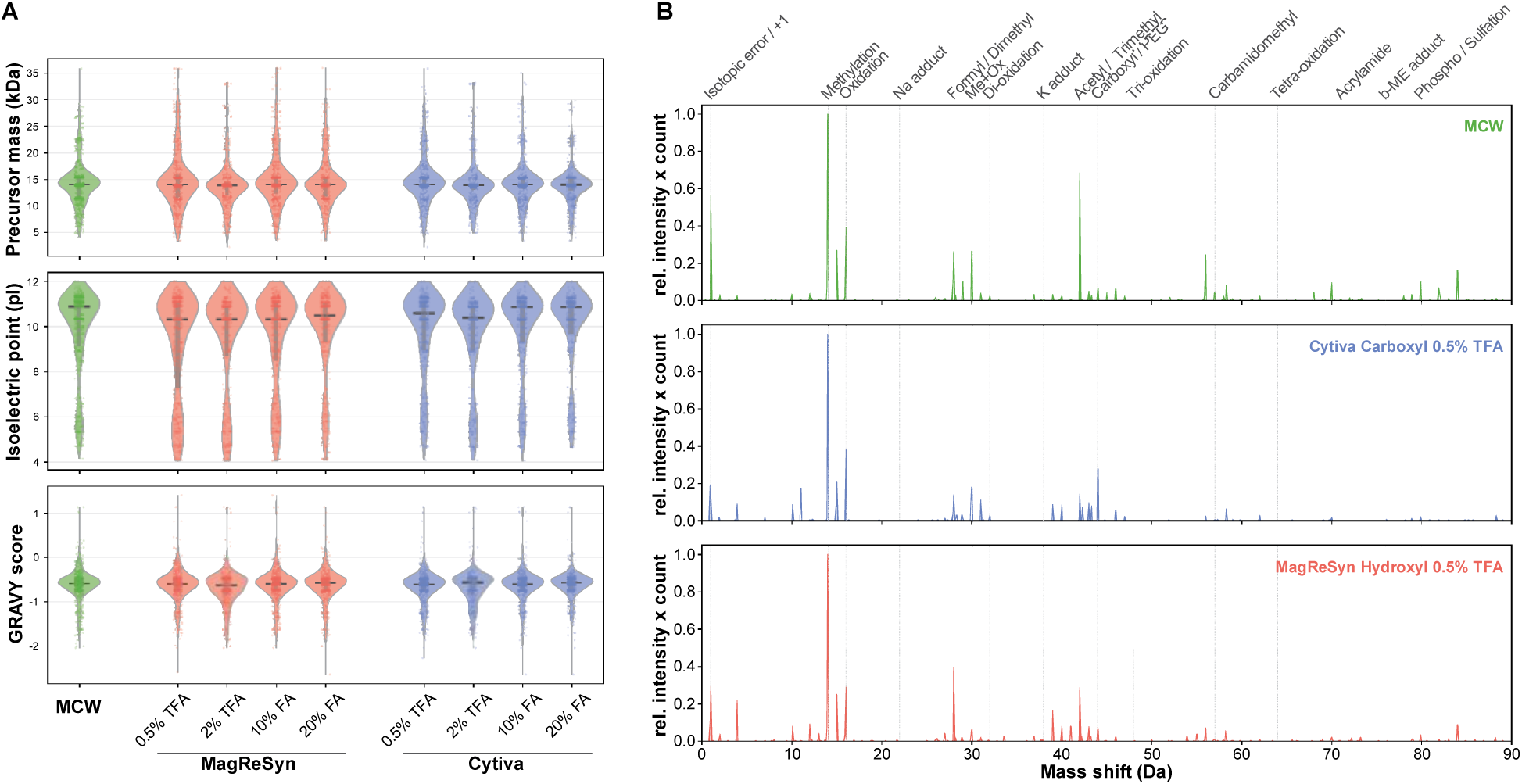
A) Violin plot of GRAVY score, isoelectric point, and mass distributions across sample types based on protein sequence. Horizontal line represents the median, and gray vertical line represents the interquartile range with individual proteoforms overlaid as jittered points. Distributions are visually and statistically indistinguishable across conditions; all omnibus effect sizes were negligible-to-small (Kruskal–Wallis η² ≤ 0.018) and every multiple-testing-corrected pairwise difference was small (Cliff’s δ ≤ 0.25). B) Mass shift analysis with MSTopDiff of MCW (green), MagReSyn with 0.5% TFA (salmon), and Cytiva with 0.5% TFA (blue). Mass shifts corresponding to common modification are annotated at the top.

### Limitations and Future Directions

While we report a sample handling workflow that enables direct recovery of protein aggregates from the surface of magnetic beads and compatibility with SDS, this work does exemplify some of the persistent challenges in top-down proteomics. In particular, the RPLC-MS/MS conducted on the Orbitrap Exploris 480 is limited by ion decay, inefficient dissociation by HCD, and spectral overlap among resulting fragment ions.^52^ Alternative acquisition modes that leverage individual ion measurements,^53,54^ ion mobility separation,^55^ additional dissociation methods,^56^ and deconvolution of overlapping species with gas-phase reactions^57,58^ would significantly improve the depth of proteoform characterization. The scope of magnetic beads and binding conditions explored here is also limited. The Cytiva and MagReSyn beads used in this study were selected due to their popularity among the proteomics community, but other commercial products or bespoke particles may outperform these beads. In addition, a more thorough examination of additives present during protein binding, such as lyotrope species, would likely yield improved capture and recovery of low abundance proteoforms.

Discussions following the initial development of SP3 have largely settled on the mechanism of protein binding as driven by protein aggregation capture (PAC) rather than mimicking hydrophilic interaction chromatography (HILIC) as initially theorized.^3,46^ The basis of PAC is that protein aggregation on beads is not dependent on bead surface type, rather that it can occur on virtually any type of beads without drastic changes in protein recovery. In contrast, the binding mechanism for HILIC relies on the presence of a hydrophilic bead surface to create a water layer for protein binding to occur. We demonstrate in this study that multiple factors attributing to both HILIC and PAC can impact protein recovery, as seen in the impact of bead construction, surface chemistry, and recovery buffer on protein elution. The beads tested here have vastly different core structures in addition to their surface functionality. Cytiva beads consist of a polystyrene core with layers of magnetite and polymer, while MagReSyn beads are constructed with a dendritic polymer surrounding a magnetite core. Combining these differences with various recovery buffers and concentrations, more research would be necessary to determine sole contributions of each factor towards protein recovery. Further, we did not extensively evaluate the impact of organic solvent choice or concentration during protein aggregation. We plan to address these limitations in future studies on this method.

We believe this approach to bead-assisted sample preparation will directly benefit practitioners of top-down, particularly in sample-limited studies. This approach is particularly well positioned for targeted TDP experiments involving immunoprecipitation as the identification of MS-compatible conditions that disrupt antigen-antibody engagement is a significant bottleneck in the development of targeted assays. SDS is nearly universally capable of releasing antigens, yet its removal is typically not possible at this stage. While SDS removal is possible by MCW precipitation, the small input protein amounts in this context (sub-microgram) leads to an unobservable protein pellet, making separation, washing, and resuspension of such very difficult. In contrast, using beads enables visual confirmation of the protein at all times.

Robotic automation provides the scalability and reproducibility necessary for clinical bottom-up proteomics studies, and magnetic bead particles that offer affinity enrichment or sample cleanup are an enabling component.^59,60^ Despite its application for biomarker discovery in modestly sized clinical cohorts,^6,9,7^ top-down proteomics has not achieved similar levels of automation. Online microfluidic sample preparation for flow injection has been enabling for native and charge-detection mass spectrometry,^25,61,62,27,29^ but a similar extent of automation has not been extended to studies requiring fractionation and LC separation. Recent work has shown that selective formation of protein coronas on nanoparticles enhanced the depth of TDP analysis of plasma by 4-fold.^63^ While this study still included PEPPI fractionation, the results therein and our findings provide optimism for a bead-based automated workflow that can support clinical TDP studies.

## Conclusions

We have developed an SP3-based approach that offers direct TDP sample clean-up. By aggregating proteins onto magnetic beads using organic solvent, SP3 enables both detergent removal and minimal sample handling, since the proteins remain associated with the bead matrix throughout the workflow until the final elution step. This single-pot format effectively reduces transfer-related losses and preserves a greater fraction of low-abundance species. A key observation was the impact that bead surface chemistry has on protein resolubilization and recovery. By combining bead-mediated aggregation with controlled solvent conditions and appropriate elution buffers, this workflow addresses the dual challenge of detergent removal and proteoform preservation, offering a solution for preparing intact protein samples compatible with top-down mass spectrometry.

## ASSOCIATED CONTENT

### Data Availability Statement

LC-MS data files have been deposited to ProteomeXchange Consortium via the MassIVE partner repository with the dataset identifier PXD080383. Source code for data analysis and figure generation is available at https://github.com/drown-lab/pac-cleanup-tdms.

### Supporting Information

The following files are available free of charge. Supplementary materials, methods, and figures (.pdf)

Table of all proteoform hits across lysate samples (.xlsx)

## AUTHOR INFORMATION

### Author Contributions

The experiments were performed by I.F. and B.D. Data analysis and manuscript writing were performed by I.F. and B.D. All authors have given approval to the final version of the manuscript.

### Notes

The authors declare no competing financial interest(s).

## ACKNOWLEDGMENT

The authors would like to thank Maria Marginteau and Miranda Muhoberac for their advice in developing this manuscript. Claude Opus 4.8 was used in drafting code.

## ABBREVIATIONS

ACN: acetonitrile
AGC: automatic gain control
AmBic: ammonium bicarbonate
BSA: bovine serum albumin
BUP: bottom-up proteomics
DMEM: Dulbecco’s Modified Eagle Medium
DTT: dithiothreitol
ESI: electrospray ionization
GRAVY: grand average of hydropathy
HCD: higher-energy collisional dissociation
HEPES: 4-(2-hydroxyethyl)-1-piperazineethanesulfonic acid
HILIC: hydrophilic interaction chromatography
HPLC: high-performance liquid chromatography
IQR: interquartile range
LC-MS/MS: liquid chromatography–tandem mass spectrometry
LOD: limit of detection
MCW: methanol–chloroform–water
MS: mass spectrometry
MS/MS: tandem mass spectrometry
*m/z*: mass-to-charge ratio
NRTDP: National Resource for Translational and Developmental Proteomics
PAC: protein aggregation capture
PEPPI: passively eluting proteins from polyacrylamide gels as intact species
pI: isoelectric point
PTM: post-translational modification
RPLC: reversed-phase liquid chromatography
FTRP: Fourier transform resolving power
S/N: signal-to-noise ratio
SDS: sodium dodecyl sulfate
SP3: single-pot, solid-phase-enhanced sample preparation
TDP: top-down proteomics
TFA: trifluoroacetic acid
TIC: total ion chromatogram.

## REFERENCES

(1) Donnelly, D. P.; Rawlins, C. M.; DeHart, C. J.; Fornelli, L.; Schachner, L. F.; Lin, Z.; Lippens, J. L.; Aluri, K. C.; Sarin, R.; Chen, B.; Lantz, C.; Jung, W.; Johnson, K. R.; Koller, A.; Wolff, J. J.; Campuzano, I. D. G.; Auclair, J. R.; Ivanov, A. R.; Whitelegge, J. P.; Paša-Tolić, L.; Chamot-Rooke, J.; Danis, P. O.; Smith, L. M.; Tsybin, Y. O.; Loo, J. A.; Ge, Y.; Kelleher, N. L.; Agar, J. N. Best Practices and Benchmarks for Intact Protein Analysis for Top-down Mass Spectrometry. Nat. Methods 2019, 16 (7), 587–594. 10.1038/s41592-019-0457-0.

(2) Kaulich, P. T.; Jeong, K.; Kohlbacher, O.; Tholey, A. Influence of Different Sample Preparation Approaches on Proteoform Identification by Top-down Proteomics. Nat Methods 2024, 21 (12), 2397–2407. 10.1038/s41592-024-02481-6.

(3) Hughes, C. S.; Foehr, S.; Garfield, D. A.; Furlong, E. E.; Steinmetz, L. M.; Krijgsveld, J. Ultrasensitive Proteome Analysis Using Paramagnetic Bead Technology. Molecular Systems Biology 2014, 10 (10), 757. 10.15252/msb.20145625.

(4) Sielaff, M.; Kuharev, J.; Bohn, T.; Hahlbrock, J.; Bopp, T.; Tenzer, S.; Distler, U. Evaluation of FASP, SP3, and iST Protocols for Proteomic Sample Preparation in the Low Microgram Range. J. Proteome Res. 2017, 16 (11), 4060–4072. 10.1021/acs.jproteome.7b00433.

(5) Takemori, A.; Sugiyama, N.; Kline, J. T.; Fornelli, L.; Takemori, N. Gel-Based Sample Fractionation with SP3-Purification for Top-Down Proteomics. J. Proteome Res. 2025, 24 (2), 850–860. 10.1021/acs.jproteome.4c00941.

(6) Melani, R. D.; Gerbasi, V. R.; Anderson, L. C.; Sikora, J. W.; Toby, T. K.; Hutton, J. E.; Butcher, D. S.; Negrão, F.; Seckler, H. S.; Srzentić, K.; Fornelli, L.; Camarillo, J. M.; LeDuc, R. D.; Cesnik, A. J.; Lundberg, E.; Greer, J. B.; Fellers, R. T.; Robey, M. T.; DeHart, C. J.; Forte, E.; Hendrickson, C. L.; Abbatiello, S. E.; Thomas, P. M.; Kokaji, A. I.; Levitsky, J.; Kelleher, N. L. The Blood Proteoform Atlas: A Reference Map of Proteoforms in Human Hematopoietic Cells. Science 2022, 375 (6579), 411–418. 10.1126/science.aaz5284.

(7) Fulcher, J. M.; Ives, A. N.; Tasaki, S.; Kelly, S. S.; Williams, S. M.; Fillmore, T. L.; Zhou, M.; Moore, R. J.; Qian, W.-J.; Paša-Tolić, L.; Yu, L.; Oveisgharan, S.; Bennett, D. A.; De Jager, P. L.; Petyuk, V. A. Discovery of Proteoforms Associated With Alzheimer’s Disease Through Quantitative Top-Down Proteomics. Molecular & Cellular Proteomics 2025, 24 (6), 100983. 10.1016/j.mcpro.2025.100983.

(8) Chapman, E. A.; Aballo, T. J.; Melby, J. A.; Zhou, T.; Price, S. J.; Rossler, K. J.; Lei, I.; Tang, P. C.; Ge, Y. Defining the Sarcomeric Proteoform Landscape in Ischemic Cardiomyopathy by Top-Down Proteomics. J. Proteome Res. 2023, 22 (3), 931–941. 10.1021/acs.jproteome.2c00729.

(9) Forte, E.; Sanders, J. M.; Pla, I.; Kanchustambham, V. L.; Hollas, M. A. R.; Huang, C.-F.; Sanchez, A.; Peterson, K. N.; Melani, R. D.; Huang, A.; Polineni, P.; Doll, J. M.; Dietch, Z.; Kelleher, N. L.; Ladner, D. P. Top-Down Proteomics Identifies Plasma Proteoform Signatures of Liver Cirrhosis Progression. Molecular & Cellular Proteomics 2024, 23 (12). 10.1016/j.mcpro.2024.100876.

(10) Ntai, I.; Fornelli, L.; DeHart, C. J.; Hutton, J. E.; Doubleday, P. F.; LeDuc, R. D.; van Nispen, A. J.; Fellers, R. T.; Whiteley, G.; Boja, E. S.; Rodriguez, H.; Kelleher, N. L. Precise Characterization of KRAS4b Proteoforms in Human Colorectal Cells and Tumors Reveals Mutation/Modification Cross-Talk. Proceedings of the National Academy of Sciences 2018, 115 (16), 201716122. 10.1073/pnas.1716122115.

(11) Adams, L. M.; DeHart, C. J.; Drown, B. S.; Anderson, L. C.; Bocik, W.; Boja, E. S.; Hiltke, T. M.; Hendrickson, C. L.; Rodriguez, H.; Caldwell, M.; Vafabakhsh, R.; Kelleher, N. L. Mapping the KRAS Proteoform Landscape in Colorectal Cancer Identifies Truncated KRAS4B That Decreases MAPK Signaling. J. Biol. Chem. 2022, 299 (1), 102768. 10.1016/j.jbc.2022.102768.

(12) D’Ippolito, R. A.; Scheidemantle, G. M.; Smith, B. P.; Powell, K.; Eury, S.; Neish, A.; Mehalko, J.; Beaumont, L.; Fer, N.; Wall, V.; Burgan, W.; Maciag, A. E.; Esposito, D.; DeHart, C. J. FLAG-KRAS4B as a Model System for KRAS4B Proteoform and PTM Evaluation by Mass Spectrometry. In KRAS: Methods and Protocols; Stephen, A. G., Esposito, D., Eds.; Springer US: New York, NY, 2024; pp 299–322. 10.1007/978-1-0716-3822-4_22.

(13) Killinger, B. A.; Madaj, Z.; Sikora, J. W.; Rey, N.; Haas, A. J.; Vepa, Y.; Lindqvist, D.; Chen, H.; Thomas, P. M.; Brundin, P.; Brundin, L.; Labrie, V. The Vermiform Appendix Impacts the Risk of Developing Parkinson’s Disease. Sci. Transl. Med. 2018, 10 (465), eaar5280. 10.1126/scitranslmed.aar5280.

(14) Chapman, E. A.; Roberts, D. S.; Tiambeng, T. N.; Andrews, J.; Wang, M.-D.; Reasoner, E. A.; Melby, J. A.; Li, B. H.; Kim, D.; Alpert, A. J.; Jin, S.; Ge, Y. Structure and Dynamics of Endogenous Cardiac Troponin Complex in Human Heart Tissue Captured by Native Nanoproteomics. Nat Commun 2023, 14 (1), 8400. 10.1038/s41467-023-43321-z.

(15) Seckler, H. D. S.; Fornelli, L.; Mutharasan, R. K.; Thaxton, C. S.; Fellers, R.; Daviglus, M.; Sniderman, A.; Rader, D.; Kelleher, N. L.; Lloyd-Jones, D. M.; Compton, P. D.; Wilkins, J. T. A Targeted, Differential Top-Down Proteomic Methodology for Comparison of ApoA-I Proteoforms in Individuals with High and Low HDL Efflux Capacity. J. Proteome Res. 2018, 17 (6), 2156–2164. 10.1021/acs.jproteome.8b00100.

(16) Wilkins, J. T.; Seckler, H. S.; Rink, J.; Compton, P. D.; Fornelli, L.; Thaxton, C. S.; LeDuc, R.; Jacobs, D.; Doubleday, P. F.; Sniderman, A.; Lloyd-Jones, D. M.; Kelleher, N. L. Spectrum of Apolipoprotein AI and Apolipoprotein AII Proteoforms and Their Associations With Indices of Cardiometabolic Health: The CARDIA Study. J. Am. Heart Assoc. 2021, 10 (17), e019890. 10.1161/JAHA.120.019890.

(17) Vilà-Rico, M.; Colomé-Calls, N.; Martín-Castel, L.; Gay, M.; Azorín, S.; Vilaseca, M.; Planas, A.; Canals, F. Quantitative Analysis of Post-Translational Modifications in Human Serum Transthyretin Associated with Familial Amyloidotic Polyneuropathy by Targeted LC–MS and Intact Protein MS. Journal of Proteomics 2015, 127, 234–246. 10.1016/j.jprot.2015.04.016.

(18) den Boer, M. A.; Greisch, J.-F.; Tamara, S.; Bondt, A.; Heck, A. J. R. Selectivity over Coverage in de Novo Sequencing of IgGs. Chem. Sci. 2020, 11 (43), 11886–11896. 10.1039/D0SC03438J.

(19) Bondt, A.; Hoek, M.; Tamara, S.; de Graaf, B.; Peng, W.; Schulte, D.; van Rijswijck, D. M. H.; den Boer, M. A.; Greisch, J.-F.; Varkila, M. R. J.; Snijder, J.; Cremer, O. L.; Bonten, M. J. M.; Heck, A. J. R. Human Plasma IgG1 Repertoires Are Simple, Unique, and Dynamic. cels 2021, 12 (12), 1131–1143.e5. 10.1016/j.cels.2021.08.008.

(20) de Graaf, S. C.; Hoek, M.; Tamara, S.; Heck, A. J. R. A Perspective toward Mass Spectrometry-Based de Novo Sequencing of Endogenous Antibodies. MAbs 2022, 14 (1), 2079449. 10.1080/19420862.2022.2079449.

(21) van Rijswijck, D. M. H.; Bondt, A.; Raafat, D.; Holtfreter, S.; Wietschel, K. A.; van der Lans, S. P. A.; Völker, U.; Bröker, B. M.; Heck, A. J. R. Persistent IgG1 Clones Dominate and Personalize the Plasma Antibody Repertoire. Science Advances 2025, 11 (16), eadt7746. 10.1126/sciadv.adt7746.

(22) Kafader, J. O.; Beu, S. C.; Early, B. P.; Melani, R. D.; Durbin, K. R.; Zabrouskov, V.; Makarov, A. A.; Maze, J. T.; Shinholt, D. L.; Yip, P. F.; Kelleher, N. L.; Compton, P. D.; Senko, M. W. STORI Plots Enable Accurate Tracking of Individual Ion Signals. J. Am. Soc. Mass Spectrom. 2019, 30 (11), 2200–2203. 10.1007/s13361-019-02309-0.

(23) Drown, B. S.; Gupta, R.; McGee, J. P.; Hollas, M. A. R.; Hergenrother, P. J.; Kafader, J. O.; Kelleher, N. L. Precise Readout of MEK1 Proteoforms upon MAPK Pathway Modulation by Individual Ion Mass Spectrometry. Anal. Chem. 2024, 96 (11), 4455–4462. 10.1021/acs.analchem.3c04758.

(24) Lu, D.; Faizi, M.; Drown, B.; Simerzin, A.; François, J.; Bradshaw, G.; Kelleher, N.; Jambhekar, A.; Gunawardena, J.; Lahav, G. Temporal Regulation of Gene Expression through Integration of P53 Dynamics and Modifications. Science Advances 2024, 10 (43), eadp2229. 10.1126/sciadv.adp2229.

(25) Xu, T.; Su, T.; Soye, B. J. D.; Kandi, S.; Huang, C.-F.; Wilkins, J. T.; Castellani, R. J.; Kafader, J. O.; Patrie, S. M.; Vassar, R.; Kelleher, N. L. The Proteoform Landscape of Tau from the Human Brain. J. Proteome Res. 2025, 24 (6), 2916–2925. 10.1021/acs.jproteome.5c00139.

(26) Huang, C.-F.; Su, T.; Flozak, A. S.; Gottardi, C. J.; Kelleher, N. L. Intact Mass Profiling Reveals Phospho-Proteoforms of the Catenins (85–110 kDa) Regulated by Actomyosin Contractility. Angewandte Chemie International Edition 2025, 64 (52), e18593. 10.1002/anie.202518593.

(27) Melani, R. D.; Des Soye, B. J.; Kafader, J. O.; Forte, E.; Hollas, M.; Blagojevic, V.; Negrão, F.; McGee, J. P.; Drown, B.; Lloyd-Jones, C.; Seckler, H. S.; Camarillo, J. M.; Compton, P. D.; LeDuc, R. D.; Early, B.; Fellers, R. T.; Cho, B.-K.; Mattamana, B. B.; Goo, Y. A.; Thomas, P. M.; Ash, M. K.; Bhimalli, P. P.; Al-Harthi, L.; Sha, B. E.; Schneider, J. R.; Kelleher, N. L. Next-Generation Serology by Mass Spectrometry: Readout of the SARS-CoV-2 Antibody Repertoire. J. Proteome Res. 2022, 21 (1), 274–288. 10.1021/acs.jproteome.1c00882.

(28) Forte, E.; Des Soye, B. J.; Melani, R. D.; Hollas, M. A. R.; Kafader, J. O.; Sha, B. E.; Schneider, J. R.; Kelleher, N. L. Divergent Antibody Repertoires Found for Omicron versus Wuhan SARS-CoV-2 Strains Using Ig-MS. J. Proteome Res. 2022, 21 (12), 2987–2997. 10.1021/acs.jproteome.2c00514.

(29) Des Soye, B. J.; McGee, J. P.; Hollas, M. A. R.; Forte, E.; Fellers, R. T.; Melani, R. D.; Wilkins, J. T.; Compton, P. D.; Kafader, J. O.; Kelleher, N. L. Automated Immunoprecipitation, Sample Preparation, and Individual Ion Mass Spectrometry Platform for Proteoforms. Anal. Chem. 2024, 96 (34), 13879–13887. 10.1021/acs.analchem.4c01962.

(30) Des Soye, B. J.; Melani, R. D.; Hollas, M. A. R.; Duan, J.; Patrie, S. M.; Fisher, T. D.; Mattamana, B. B.; Daud, A.; Pinelli, D. F.; Ladner, D. P.; Kelleher, N. L.; Forte, E. Characterization of the Antibody Response to SARS-CoV-2 Infection in COVID-19 Transplant versus Nontransplant Recipients by Ig-MS. J. Proteome Res. 2024, 23 (9), 3944–3957. 10.1021/acs.jproteome.4c00285.

(31) Kachuk, C.; Doucette, A. A. The Benefits (and Misfortunes) of SDS in Top-down Proteomics. Journal of Proteomics 2018, 175, 75–86. 10.1016/j.jprot.2017.03.002.

(32) Bligh, E. G.; Dyer, W. J. A Rapid Method of Total Lipid Extraction and Purification. Can J Biochem Physiol 1959, 37 (8), 911–917. 10.1139/o59-099.

(33) Wessel, D.; Flügge, U. I. A Method for the Quantitative Recovery of Protein in Dilute Solution in the Presence of Detergents and Lipids. Anal Biochem 1984, 138 (1), 141–143. 10.1016/0003-2697(84)90782-6.

(34) le Maire, M.; Deschamps, S.; Møller, J. V.; Le Caer, J. P.; Rossier, J. Electrospray Ionization Mass Spectrometry on Hydrophobic Peptides Electroeluted from Sodium Dodecyl Sulfate-Polyacrylamide Gel Electrophoresis Application to the Topology of the Sarcoplasmic Reticulum Ca2+ ATPase. Anal Biochem 1993, 214 (1), 50–57. 10.1006/abio.1993.1455.

(35) Puchades, M.; Westman, A.; Blennow, K.; Davidsson, P. Removal of Sodium Dodecyl Sulfate from Protein Samples Prior to Matrix-Assisted Laser Desorption/Ionization Mass Spectrometry. Rapid Commun. Mass Spectrom. 1999, 13 (5), 344–349. 10.1002/(SICI)1097-0231(19990315)13:5%3C344::AID-RCM489%3E3.0.CO;2-V.

(36) Takemori, A.; Kaulich, P. T.; Cassidy, L.; Takemori, N.; Tholey, A. Size-Based Proteome Fractionation through Polyacrylamide Gel Electrophoresis Combined with LC–FAIMS–MS for In-Depth Top-Down Proteomics. Anal. Chem. 2022, 94 (37), 12815–12821. 10.1021/acs.analchem.2c02777.

(37) Tran, J. C.; Doucette, A. A. Gel-Eluted Liquid Fraction Entrapment Electrophoresis: An Electrophoretic Method for Broad Molecular Weight Range Proteome Separation. Anal. Chem. 2008, 80 (5), 1568–1573. 10.1021/ac702197w.

(38) Takemori, A.; Butcher, D. S.; Harman, V. M.; Brownridge, P.; Shima, K.; Higo, D.; Ishizaki, J.; Hasegawa, H.; Suzuki, J.; Yamashita, M.; Loo, J. A.; Loo, R. R. O.; Beynon, R. J.; Anderson, L. C.; Takemori, N. PEPPI-MS: Polyacrylamide-Gel-Based Prefractionation for Analysis of Intact Proteoforms and Protein Complexes by Mass Spectrometry. J. Proteome Res. 2020, 19 (9), 3779–3791. 10.1021/acs.jproteome.0c00303.

(39) Cai, W.; Tucholski, T.; Chen, B.; Alpert, A. J.; McIlwain, S.; Kohmoto, T.; Jin, S.; Ge, Y. Top-Down Proteomics of Large Proteins up to 223 kDa Enabled by Serial Size Exclusion Chromatography Strategy. Anal. Chem. 2017, 89 (10), 5467–5475. 10.1021/acs.analchem.7b00380.

(40) Yang, Z.; Shen, X.; Chen, D.; Sun, L. Toward a Universal Sample Preparation Method for Denaturing Top-Down Proteomics of Complex Proteomes. J. Proteome Res. 2020, 19 (8), 3315–3325. 10.1021/acs.jproteome.0c00226.

(41) Cassidy, L.; Helbig, A. O.; Kaulich, P. T.; Weidenbach, K.; Schmitz, R. A.; Tholey, A. Multidimensional Separation Schemes Enhance the Identification and Molecular Characterization of Low Molecular Weight Proteomes and Short Open Reading Frame-Encoded Peptides in Top-down Proteomics. J Proteomics 2021, 230, 103988. 10.1016/j.jprot.2020.103988.

(42) Meng, F.; Cargile, B. J.; Patrie, S. M.; Johnson, J. R.; McLoughlin, S. M.; Kelleher, N. L. Processing Complex Mixtures of Intact Proteins for Direct Analysis by Mass Spectrometry. Anal. Chem. 2002, 74 (13), 2923–2929. 10.1021/ac020049i.

(43) Brown, K. A.; Chen, B.; Guardado-Alvarez, T. M.; Lin, Z.; Hwang, L.; Ayaz-Guner, S.; Jin, S.; Ge, Y. A Photocleavable Surfactant for Top-down Proteomics. Nat Methods 2019, 16 (5), 417–420. 10.1038/s41592-019-0391-1.

(44) Laganowsky, A.; Reading, E.; Hopper, J. T. S.; Robinson, C. V. Mass Spectrometry of Intact Membrane Protein Complexes. Nat Protoc 2013, 8 (4), 639–651. 10.1038/nprot.2013.024.

(45) Hughes, C. S.; Moggridge, S.; Müller, T.; Sorensen, P. H.; Morin, G. B.; Krijgsveld, J. Single-Pot, Solid-Phase-Enhanced Sample Preparation for Proteomics Experiments. Nat Protoc 2019, 14 (1), 68–85. 10.1038/s41596-018-0082-x.

(46) Batth, T. S.; Tollenaere, M. A. X.; Rüther, P.; Gonzalez-Franquesa, A.; Prabhakar, B. S.; Bekker-Jensen, S.; Deshmukh, A. S.; Olsen, J. V. Protein Aggregation Capture on Microparticles Enables Multipurpose Proteomics Sample Preparation. Mol. Cell. Proteom. 2019, 18 (5), 1027–1035. 10.1074/mcp.TIR118.001270.

(47) Dagley, L. F.; Infusini, G.; Larsen, R. H.; Sandow, J. J.; Webb, A. I. Universal Solid-Phase Protein Preparation (USP3) for Bottom-up and Top-down Proteomics. J. Proteome Res. 2019, 18 (7), 2915–2924. 10.1021/acs.jproteome.9b00217.

(48) Smeriglio, N.; Li, H.; Mazli, W. N. A. B.; Bendel, K.; Hao, L. Contaminant Spot Check and Removal Assay (ContamSPOT) for Mass Spectrometry Analysis. Anal. Chem. 2024, 96 (6), 2574–2581. 10.1021/acs.analchem.3c05020.

(49) LeDuc, R. D.; Fellers, R. T.; Early, B. P.; Greer, J. B.; Thomas, P. M.; Kelleher, N. L. The C-Score: A Bayesian Framework to Sharply Improve Proteoform Scoring in High-Throughput Top down Proteomics. J. Proteome Res. 2014, 13 (7), 3231–3240. 10.1021/pr401277r.

(50) Jeong, K.; Kim, J.; Gaikwad, M.; Hidayah, S. N.; Heikaus, L.; Schlüter, H.; Kohlbacher, O. FLASHDeconv: Ultrafast, High-Quality Feature Deconvolution for Top-Down Proteomics. Cell Systems 2020, 10 (2), 213–218.e6. 10.1016/j.cels.2020.01.003.

(51) Feroz, A.; Nagornov, K. O.; Müller, T.; Sachsenberg, T.; Tsybin, Y.; Kohlbacher, O.; Jeong, K. False Discovery Rate Estimation in Spectral Deconvolution in Top-down Proteomics. Authorea 2025 (0527). 10.22541/au.174834556.69764899/v1.

(52) Compton, P. D.; Zamdborg, L.; Thomas, P. M.; Kelleher, N. L. On the Scalability and Requirements of Whole Protein Mass Spectrometry. Anal. Chem. 2011, 83 (17), 6868–6874. 10.1021/ac2010795.

(53) Kafader, J. O.; Melani, R. D.; Durbin, K. R.; Ikwuagwu, B.; Early, B. P.; Fellers, R. T.; Beu, S. C.; Zabrouskov, V.; Makarov, A. A.; Maze, J. T.; Shinholt, D. L.; Yip, P. F.; Tullman-Ercek, D.; Senko, M. W.; Compton, P. D.; Kelleher, N. L. Multiplexed Mass Spectrometry of Individual Ions Improves Measurement of Proteoforms and Their Complexes. Nat. Methods 2020, 17 (4), 391–394. 10.1038/s41592-020-0764-5.

(54) Kafader, J. O.; Durbin, K. R.; Melani, R. D.; Soye, B. J. D.; Schachner, L. F.; Senko, M. W.; Compton, P. D.; Kelleher, N. L. Individual Ion Mass Spectrometry Enhances the Sensitivity and Sequence Coverage of Top-Down Mass Spectrometry. J. Proteome Res. 2020, 19 (3), 1346–1350. 10.1021/acs.jproteome.9b00797.

(55) Gerbasi, V. R.; Melani, R. D.; Abbatiello, S. E.; Belford, M. W.; Huguet, R.; McGee, J. P.; Dayhoff, D.; Thomas, P. M.; Kelleher, N. L. Deeper Protein Identification Using Field Asymmetric Ion Mobility Spectrometry in Top-Down Proteomics. Anal. Chem. 2021, 93 (16), 6323–6328. 10.1021/acs.analchem.1c00402.

(56) Shaw, J. B.; Li, W.; Holden, D. D.; Zhang, Y.; Griep-Raming, J.; Fellers, R. T.; Early, B. P.; Thomas, P. M.; Kelleher, N. L.; Brodbelt, J. S. Complete Protein Characterization Using Top-Down Mass Spectrometry and Ultraviolet Photodissociation. J Am Chem Soc 2013, 135 (34), 12646–12651. 10.1021/ja4029654.

(57) McLuckey, S. A.; Stephenson, J. L.; Asano, K. G. Ion/Ion Proton-Transfer Kinetics: Implications for Analysis of Ions Derived from Electrospray of Protein Mixtures. Anal. Chem. 1998, 70 (6), 1198–1202. 10.1021/ac9710137.

(58) Huguet, R.; Mullen, C.; Srzentić, K.; Greer, J. B.; Fellers, R. T.; Zabrouskov, V.; Syka, J. E. P.; Kelleher, N. L.; Fornelli, L. Proton Transfer Charge Reduction Enables High-Throughput Top-Down Analysis of Large Proteoforms. Anal. Chem. 2019, 91 (24), 15732–15739. 10.1021/acs.analchem.9b03925.

(59) Kverneland, A. H.; Harking, F.; Vej-Nielsen, J. M.; Huusfeldt, M.; Bekker-Jensen, D. B.; Svane, I. M.; Bache, N.; Olsen, J. V. Fully Automated Workflow for Integrated Sample Digestion and Evotip Loading Enabling High-Throughput Clinical Proteomics. Molecular & Cellular Proteomics 2024, 23 (7). 10.1016/j.mcpro.2024.100790.

(60) Schär, S.; Räss, L.; Malinovska, L.; Savickas, S.; Cavallo, F.; Below, C.; Tognetti, M.; Shichkova, P.; Gourdet, B.; Robles, G.; Iu, L.; Vowinckel, J.; Feng, Y.; Hjerpe, R.; Bruderer, R.; Reiter, L. A Flexible End-to-End Automated Sample Preparation Workflow Enables Standardized and Scalable Bottom-up Proteomics. Anal. Chem. 2025, 97 (40), 22116–22131. 10.1021/acs.analchem.5c03829.

(61) Kim, K. H.; Compton, P. D.; Tran, J. C.; Kelleher, N. L. Online Matrix Removal Platform for Coupling Gel-Based Separations to Whole Protein Electrospray Ionization Mass Spectrometry. J Proteome Res 2015, 14 (5), 2199–2206. 10.1021/pr501331q.

(62) Chilmonczyk, M. A.; Kottke, P. A.; Stevens, H. Y.; Guldberg, R. E.; Fedorov, A. G. Dynamic Mass Spectrometry Probe for Electrospray Ionization Mass Spectrometry Monitoring of Bioreactors for Therapeutic Cell Manufacturing. Biotechnology and Bioengineering 2019, 116 (1), 121–131. 10.1002/bit.26832.

(63) Huang, C.-F.; Hollas, M. A.; Sanchez, A.; Bhattacharya, M.; Ho, G.; Sundaresan, A.; Caldwell, M. A.; Zhao, X.; Benz, R.; Siddiqui, A.; Kelleher, N. L. Deep Profiling of Plasma Proteoforms with Engineered Nanoparticles for Top-Down Proteomics. J Proteome Res 2024, 23 (10), 4694–4703. 10.1021/acs.jproteome.4c00621.

