## Supplemental Information for "Protein Aggregation Capture for Top-down Proteomics"

### Bead-based Sample Preparation for Top-down Proteomics

Table of Contents

Supporting Experimental Section

Table S1: Supporting Materials and Reagents

S2: Detailed Sample Clean-up Protocols

S3: Cell Culture and Lysis Protocol

S4: Lysate Aliquoting and Clean-up

S5: ContamSpot Detergent-Removal Assay

S6: Dilute Myoglobin Experiment

S7: MSTopDiff Protocol for Mass Shift Analysis

S8: SDS-PAGE for Bead Protein Recovery

Table S2: LC-MS/MS Acquisition Parameters

Supporting Figures

Figure S1: Comparison of all proteins in NRTDP standard protein solution for MCW.

Figure S2: Comparison of all proteins in NRTDP standard protein solution for Cytiva.

Figure S3: Comparison of all proteins in NRTDP standard protein solution for MagResyn.

Figure S4: Histogram representing counts of proteoform hits across average retention time.

Figure S5: Gels for bead and recovery buffer evaluation.

Figure S6: Proteoforms across sample types and their p-score distributions with FLASHDeconv Q-scores and FDR cut-off.

Figure S7: GRAVY, isoelectric point, and mass distributions excluding histones.

Figure S8: Mass shift analysis of all cleanup conditions.

**Table S1. Materials and Reagents**

| Reagent / Consumable | Supplier | Catalog / Ref No. |
|---|---|---|
| **Pierce Intact Protein Standard Mix** | Thermo Fisher | A33527 |
| **LC-MS-grade water** | Thermo Fisher | W64 |
| **LC-MS-grade Methanol** | Fisher Scientific | A456-4 |
| **HPLC-grade Chloroform** | Sigma-Aldrich | 366927 |
| **Eppendorf protein LoBind tubes** | Fisher Scientific | 05-414-206 |
| **Glass LC-MS vials** | Fisher Scientific | 6PRV11-03FIVP |
| **MAbPac Capillary RP column (150 µm i.d. x 15 cm L)** | Thermo Fisher | 164947 |
| **C4 MicroSPE trap column** | CoAnn Technologies | SPE20005C04WF-2 |
| **DTT** | Thermo Fisher | A39255 |
| **Cytiva Sera-Mag SpeedBeads, hydrophilic** | Fisher Scientific | 09-981-121 |
| **Cytiva Sera-Mag SpeedBeads, hydrophobic** | Fisher Scientific | 09-981-123 |
| **MagReSyn Hydroxyl beads** | ReSyn Biosciences | MR-HYX005 |
| **Myoglobin (equine skeletal muscle)** | Sigma-Aldrich | M0630 |
| **Carbonic anhydrase (bovine erythrocytes)** | Sigma-Aldrich | C2624 |
| **Ubiquitin (bovine erythrocytes)** | Sigma-Aldrich | U6253 |
| **Trypsinogen (bovine pancreas)** | Sigma-Aldrich | T1143 |
| **BSA standard, 2 mg/mL** | Thermo Fisher | 23209 |
| **HEPES** | Sigma-Aldrich | H4034 |
| **NaCl** | Sigma-Aldrich | S7653 |
| **SDS** | Sigma-Aldrich | L6026 |
| **Trifluoroacetic Acid** | Fisher Scientific | A116-10X1AMP |
| **Formic Acid** | Fisher Scientific | A1110X1-AMP |
| **Acetonitrile** | Fisher Scientific | A955-1 |
| **Ethanol** | Decon Laboratories, Inc. | 04-355-223 |
| **Ammonium bicarbonate** | Supelco | 5.33005 |

| | | |
|---|---|---|
| **Halt protease inhibitor** | Thermo Scientific | PI78446 |
| **DMEM** | Corning | MT10013CV |
| **Penicillin-streptomycin** | Corning | MT30001CI |
| **Fetal Bovine Serum** | Corning | MT35016CV |
| **Dulbecco's Phosphate-Buffered Salt Solution (DPBS)** | Corning | MT21031CV |
| **LDS Sample Buffer** | Invitrogen | NP0008 |
| **2-Mercaptoethanol** | Sigma-Aldrich | M3148 |
| **PageRuler™ Plus PreStained Protein Ladder, 10 to 250 kDa** | Thermo Scientific | PI26619 |
| **Bis-Tris Plus Mini Protein Gels, 4-12%, 1.0 mm, WedgeWell™ format** | Invitrogen | NW04120BOX |
| **NuPAGE™ MES SDS Running Buffer (20X)** | Invitrogen | NP000202 |
| **Brilliant Blue G, Ultrapure** | Thermo Scientific Chemicals | AA4331809 |
| **Acetic Acid, Glacial (Certified ACS)** | Fisher Scientific | A38-500 |

#### S2. Detailed Sample Clean-up Protocols

**Methanol-chloroform-water (MCW) precipitation.** A 2.5 µL aliquot (4.5 µg) of the Pierce Intact Protein Standard was resuspended in 97.5 µL of LC-MS-grade water in a LoBind tube. Four volumes of methanol (400 µL) were added and vigorously mixed. One volume of chloroform (100 µL) was added and vigorously mixed. Three volumes of water (300 µL) were added and vigorously mixed, resulting in a cloudy mixture. The sample was centrifuged at 20,000 × g for 5 min at room temperature to induce the formation of layers. The upper layer was carefully removed, and 400 µL of methanol was layered over the interphase pellet without disturbing it. The sample was centrifuged at 20,000 × g for 10 min. The supernatant was discarded and the pellet resuspended in 50 µL of Solvent A (95% water, 5% acetonitrile, 0.2% formic acid). After a final 20,000 × g, 10 min spin, the supernatant was transferred to a glass LC-MS vial. For the NRTDP standard, a 2.5 µL aliquot with 5 µg of total protein (1:10:16:16 ratio of ubiquitin, trypsinogen, myoglobin, and carbonic anhydrase) was used and prepared the same way.

**SP3 with Cytiva carboxylate beads.** Hydrophilic and hydrophobic beads were combined 1:1, equilibrated to room temperature for 30 min, washed twice with 50 µL of water, and resuspended in water to 10 µg/µL. The Pierce aliquot (4.5 µg) was resuspended in 47.5 µL of water and reduced with DTT (10 mM final) for 30 min at 60 °C. Bead suspension (10 µL) and 60 µL of ethanol (50% final) were added, and the sample was mixed at 1000 rpm for 10 min at room temperature. Beads were collected on a magnet, washed twice with 200 µL of 80% ethanol, and proteins were eluted in 50 µL of 0.5% TFA with mixing at 1000 rpm for 20 min. The supernatant was collected into an LC-MS vial.

**SP3 with MagReSyn hydroxyl beads.** Beads were equilibrated to room temperature for 30 min, gently resuspended, transferred to 1.5 mL LoBind tube, washed twice with 50 µL of water, and resuspended in water to 10 µg/µL. The Pierce aliquot (4.5 µg) was resuspended in 47.5 µL of water and reduced with DTT (10 mM final) for 30 min at 60 °C. Bead suspension (10 µL, 100 µg) was added, followed by acetonitrile to 70% final, and the sample was mixed at 1000 rpm for 10 min at room temperature. Beads were collected on a magnet and washed twice with 200 µL of 100% acetonitrile without disrupting the bead–protein pellet. Proteins were eluted in 50 µL of 0.5% TFA with mixing at 1000 rpm for 20 min, and the supernatant was transferred to an LC-MS vial.

**S3. Cell Culture and Lysis**

HeLa cells were grown in DMEM containing L-glutamine, glucose, and sodium pyruvate, supplemented with 2% penicillin-streptomycin and 10% fetal bovine serum, at 37 °C and 5% $CO_2$, and harvested at passage 15 by scraping. Cells were washed with 10 mL of DPBS per plate, then five 15 cm plates (~100 million cells) were pelleted by centrifugation at 20,000 × g and stored at −80 °C. For lysis, cells were resuspended in 1 mL of lysis buffer (50 mM HEPES pH 8.5, 1% SDS, 1× Halt) and sonicated on ice (Fisher Scientific Model 120, 50% amplitude, 2 s on / 5 s off, 30 s total). Lysate was clarified at 10,000 × g for 30 min at 4 °C, and the supernatant was retained.

**S4. Lysate Aliquoting and Clean-up**

Clarified lysate (~2 mg in 1 mL) was reduced with DTT (10 mM final) at 60 °C for 30 min. Thirty-microliter aliquots were distributed across nine tubes: one for MCW and the remainder split between MagReSyn hydroxyl and Cytiva carboxylate bead preparations. The MCW aliquot was diluted to 100 µL with water and processed as in Section S2; besides changing the amount of recovery buffer from 50 µL to 100 µL for the final elution, bead preparations followed the corresponding S2 protocols.

**S5. ContamSpot Detergent-Removal Assay**

BSA (1.5 mg from a 2 mg/mL ampule) was added to 1.5 mL of 50 mM HEPES, 1% SDS. A 5 µL aliquot was diluted with 45 µL of 50 mM HEPES (pH 7.5), 1% SDS, mixed, reduced with 5.5 µL of 100 mM DTT (30 min, 60 °C). For MagReSyn samples, 30 µL of bead slurry was washed twice with 100 µL of water and resuspended in 50 µL of water; 25 µL was added to the BSA sample with 187.8 µL of acetonitrile (70% final), and 10 µL was reserved as the pre-cleanup ("A") sample. After 10 min mixing and magnetic collection, beads were washed three times with 100 µL of acetonitrile. Proteins were eluted with either 100 mM ammonium bicarbonate or 0.5% TFA (50 µL, 20 min mixing), and 10 µL of supernatant was reserved as the post-elution ("B") sample. For Cytiva samples, 3 µL each of hydrophobic and hydrophilic beads were washed twice with 100 µL of water, resuspended in 50 µL of water, and 25 µL added per BSA sample with ethanol to 50% final (10 µL reserved as "A"); after collection, beads were washed three times with 100 µL of ethanol and eluted as above. The MCW control was diluted to 100 µL with water and processed as in Section S2. To a 1.5 mL Eppendorf tube, 1 µL of 0.1% toluidine blue was added along with 1 µL of the sample. 3 µL of ethyl acetate was added to the tube as well, then vortexed for 15 s and centrifuged for 15 s. 1.5 µL of the ethyl acetate layer was spotted onto a TLC plate and imaged when dry. Spot intensity (O-toluidine blue) was read colorimetrically; a TFA-only reference spot was included to control for ion-pair color formation by TFA.

**S6. Dilute Myoglobin Recovery**

Equine myoglobin was dissolved and diluted to 4 ng/mL in 0.5% SDS (v/v), 50 mM HEPES (pH 7.4) and divided into 50 mL aliquots (200 ng). Each aliquot was subjected to either MCW or SP3-based cleanup (above). Protein samples were resuspended/recovered in 25 mL 2% acetonitrile, 0.1% formic acid, 97.9% water (MCW method) or 25 uL 0.5% TFA (SP3 method). Samples (5 µL injection) were analyzed with a MS1-only acquisition method.

**S7: MSTopDiff Protocol for Mass Shift Analysis**

Thermo .raw files were converted to .mzML using MSConvert (v.3.0, GUI, 64-bit), with standard options and "Peak Picking" filter and "Vendor" algorithm with standard settings. Intact mass inference was performed using FLASHDeconv (packaged with OpenMS version 3.5.0), with command line options "-write_detail -FD:report_FDR -FD:allowed_isotope_error 0 -SD:tol 10 -SD:min_mass 2000 -SD:max_mass 70000 -SD:min_charge 4 -SD:max_charge 60". Non-decoy deconvoluted masses with at least 3 charge states and q-value > 0.05 were used for mass shift analysis. Mass shifts were identified with MSTopDiff (v.1.1.0) with the settings Mass feature filter: 0-100 kDa; Retention time range: 0-60 min; Delta mass range: 0-150 Da, Retention time window: 2 minutes; Maximum charge difference: 2; Bin size: 0.01.

**S8: SDS-PAGE and Coomassie Staining Protocol**

From section S4, 10 µL of the final eluted protein sample was saved as well as the beads to the corresponding sample. The beads were washed twice with 200 µL of LC-MS grade water, then dried along with the protein samples on a CentriVap Cold Trap (Labconco, Kansas City, MO). Once dried, both the beads and proteins were resuspended in 10 µL of LDS sample buffer, 4 µL of 2-mercaptoethanol, and 12 µL of deionized water and heated at 70 ˚C for 10 minutes on a Thermal Mixer II (Fisher Scientific, Waltham, MA), with 3 µL of PageRuler Plus Prestained protein ladder. After heating, the bead samples were placed on a magnet stand, the supernatant was removed and was loaded onto a 4-12% Bis-Tris mini gel with 1 X MES running buffer and ran on an Invitrogen Mini Gel Tank (Invitrogen, Waltham, MA) at 120 V for 60 minutes. After running, the gel was stained with 0.1% Brilliant Blue G in 40% methanol and 10% acetic acid for 1 hour, then destained with 40% methanol and 10% acetic acid overnight. Gel was imaged using a ChemiDoc MP Imaging System (Bio-Rad, Hercules, CA) on the white tray with optimal auto-exposure settings.

**Table S3. LC-MS/MS Acquisition Parameters**

| Parameter | Primary method (standards, lysate) | Dilute Myoglobin |
|---|---|---|
| **Injection volume** | 2 µL | 5 µL |
| **Column** | MAbPac Capillary RP (150 mm ID x 15 cm L), direct injection | MAbPac Capillary RP (150 mm ID x 15 cm L), trap-and-elute with CoAnn $C_4$ trap (5 µm, 300A, 200 µm ID x 3 cm L) |
| **Mobile phase A / B** | 0.1% formic acid in water / 100% acetonitrile | 0.1% formic acid in water / 100% acetonitrile |

| Parameter | Primary method (standards, lysate) | Dilute Myoglobin |
|---|---|---|
| **Flow rate** | 1.5 μL/min | 1.0 μL/min |
| **Gradient** | 4%→16% B (1.5 min)→48% B (50 min)→64% B (3 min); wash/equil. 99% B, 5 min | 5% B →20% B(1.5 min) →50% B (15 min) → 80% B (10 min); wash/equil. 95% B, 3 min |
| **Instrument / mode** | Exploris 480 / intact-protein, positive, low-pressure (0.5 mTorr) | Exploris 480 / intact-protein, positive, low-pressure (0.5 mTorr) |
| **MS1 resolving power** | 120,000 | 120,000 |
| **MS1 range / AGC / µscans / IT** | 500–2000 *m/z* / 250% / 3 / 50 ms | 500–2000 *m/z* / 300% / 3 / 200 ms |
| **MS1 filters** | intensity ≥$2\times10^4$; charge 5–50; dynamic exclusion (1×, 30 s) | |
| **MS2 resolving power** | 60,000 | |
| **MS2 range / AGC / µscans** | 350–2000 *m/z* / 500% / 3 (standards), 4 (lysate) | |
| **MS2 isolation / activation / cycle** | 1.5 *m/z* isolation (standards), 2 *m/z* isolation (lysate) / 33% HCD NCE / 3 dependent scans | |

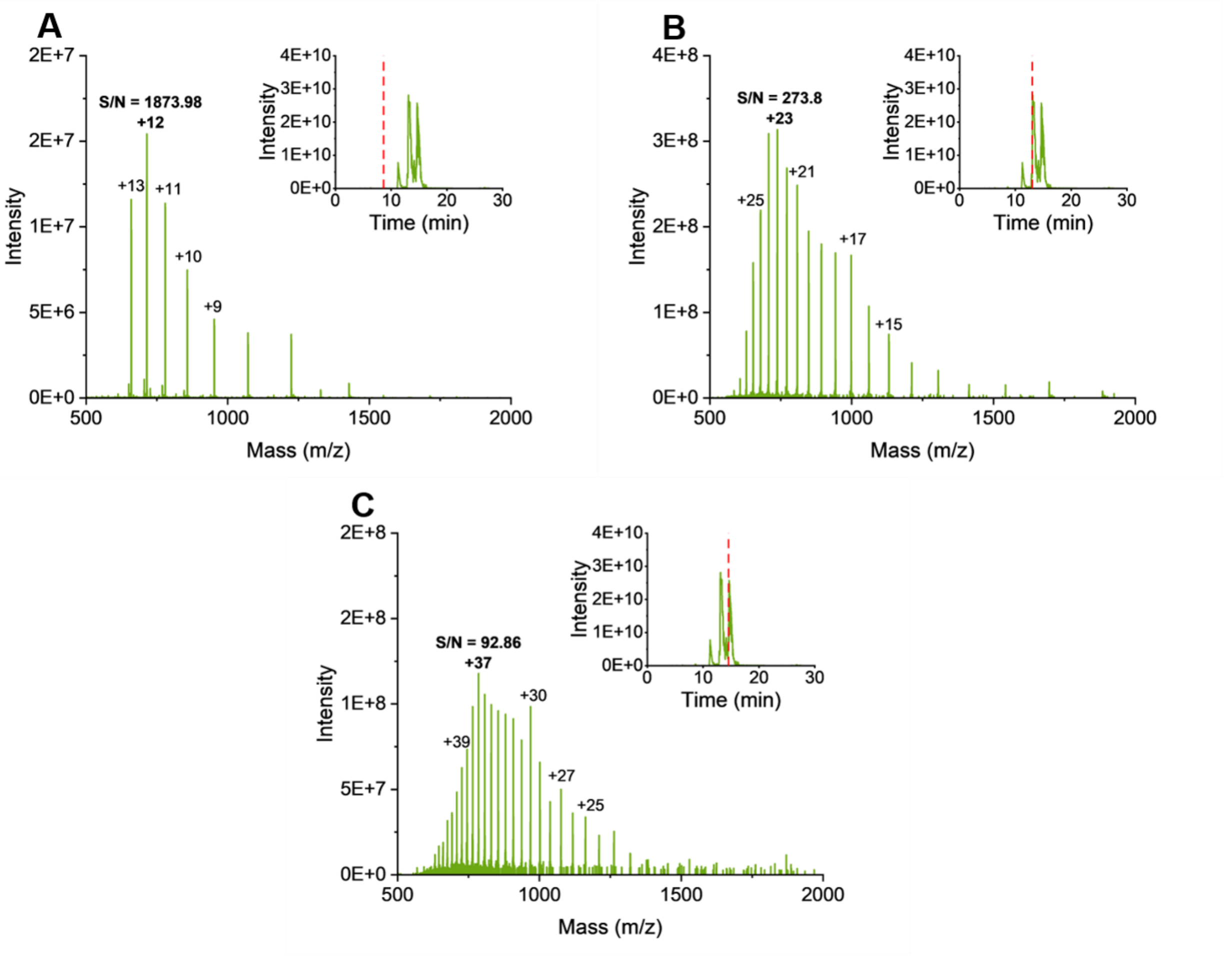


Figure S1. MS1 and location on TIC for A) ubiquitin, B) myoglobin, and C) carbonic anhydrase present in NRTDP standard for MCW sample.

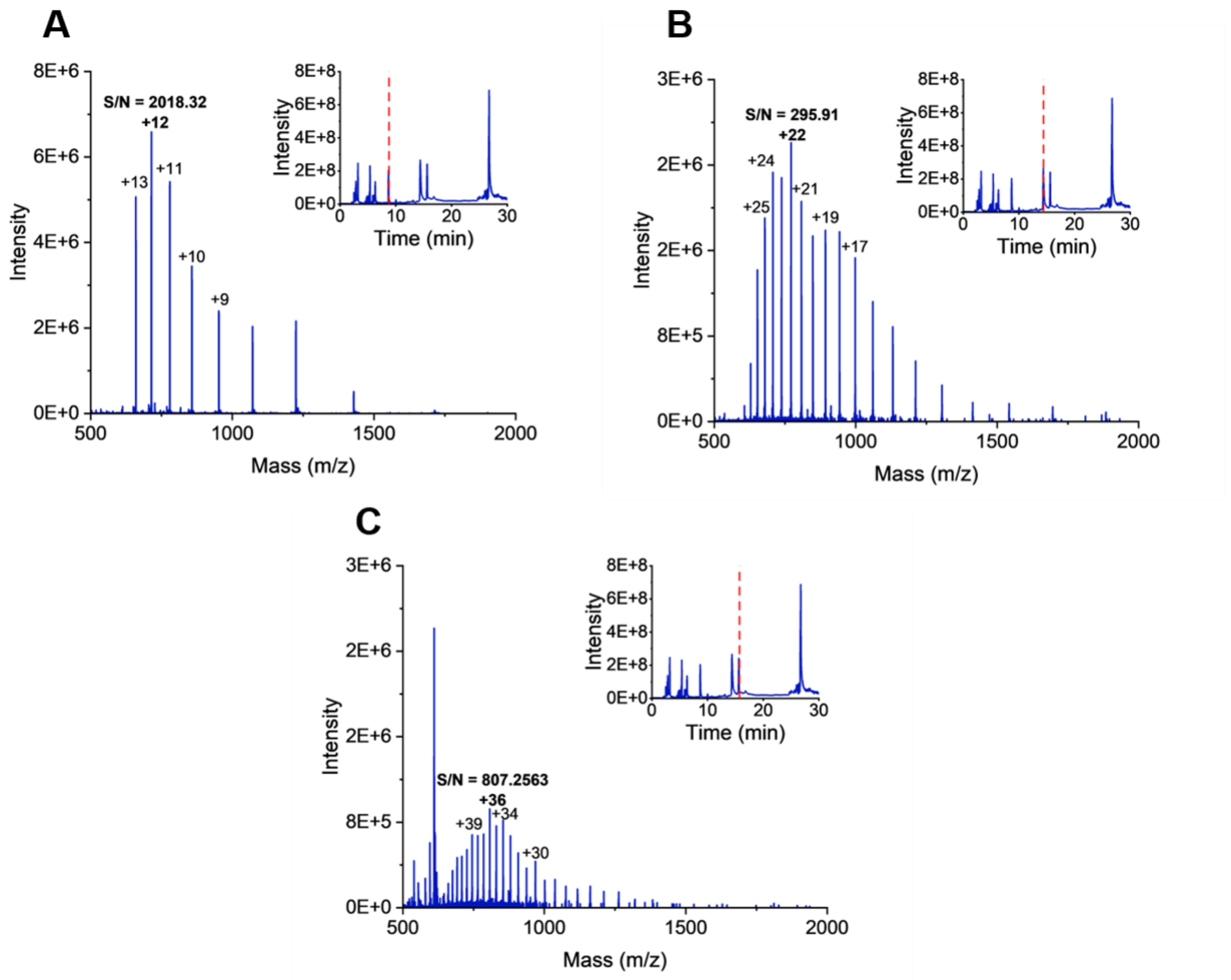


Figure S2. MS1 and location on TIC for A) ubiquitin, B) myoglobin, and C) carbonic anhydrase present in NRTDP standard for Cytiva sample.

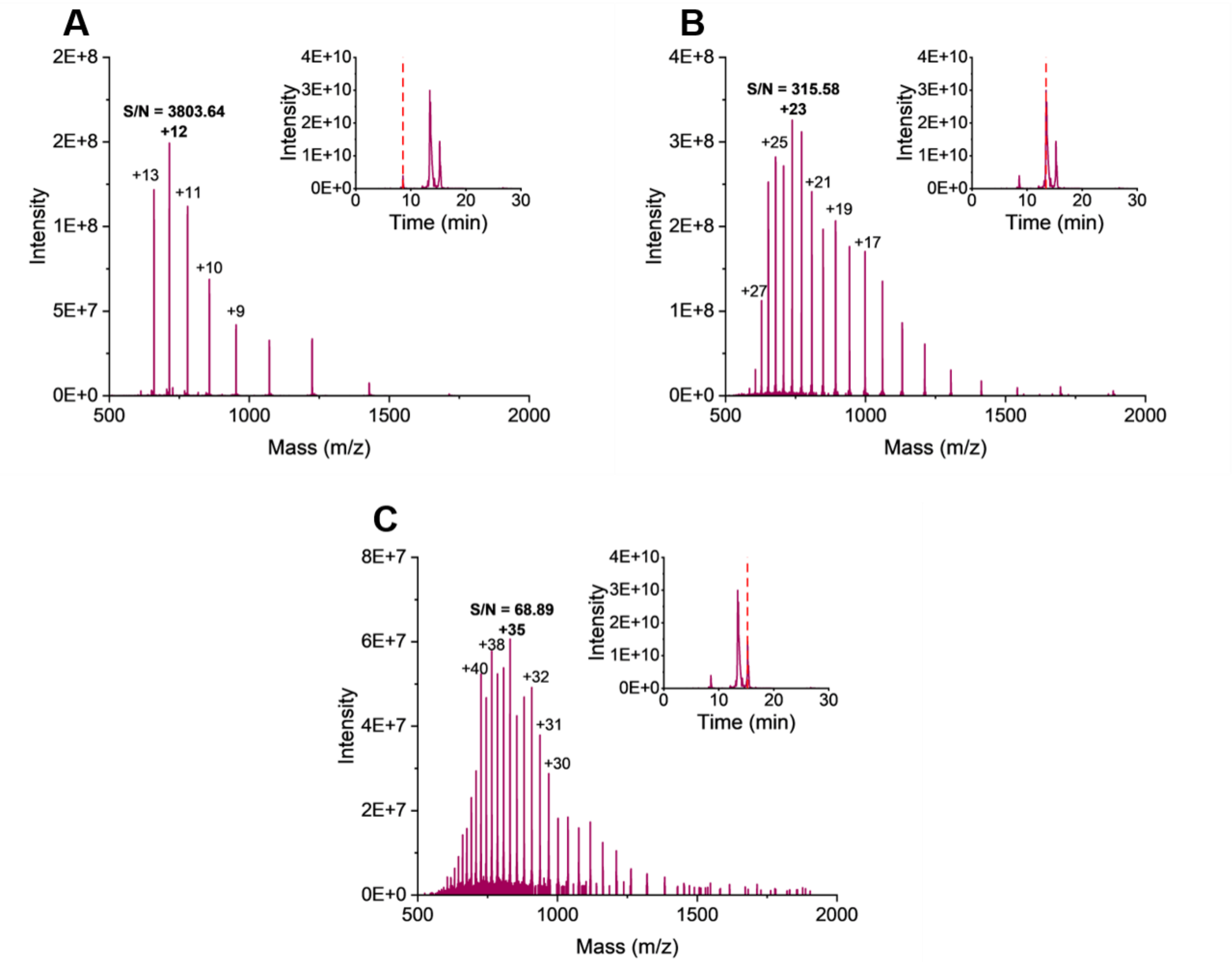


Figure S3. MS1 and location on TIC for A) ubiquitin, B) myoglobin, and C) carbonic anhydrase present in NRTDP standard for hydroxyl MagReSyn sample.

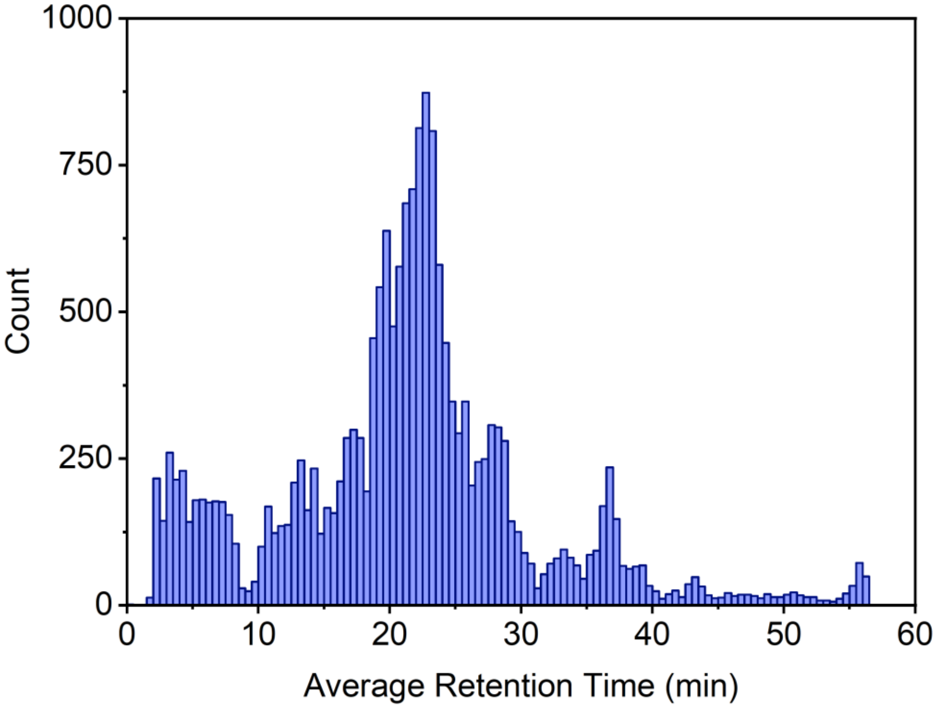


Figure S4. Retention time distribution of proteoforms for all lysate samples, bin size 0.5 (30 s).

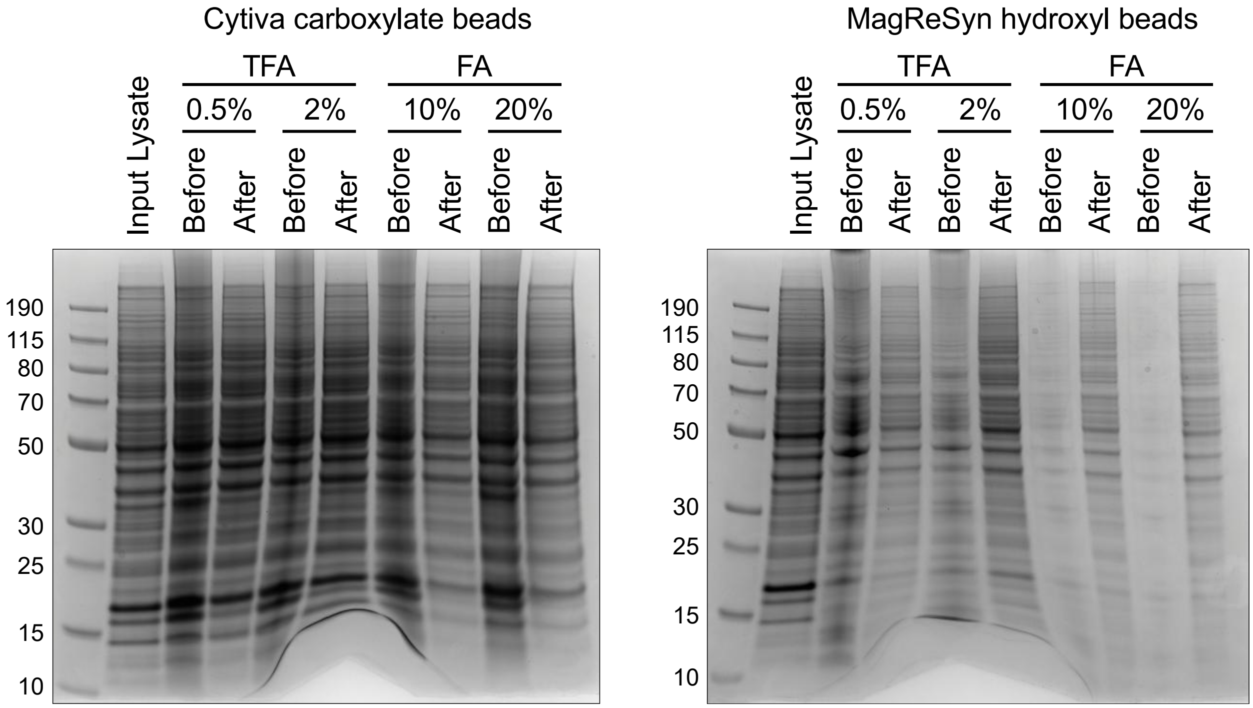

Figure S5. Coomassie stained gels for proteins that remain bound to (A) Cytiva beads and (B) MagReSyn hydroxyl beads before and after exposure to different recovery buffers and strengths.

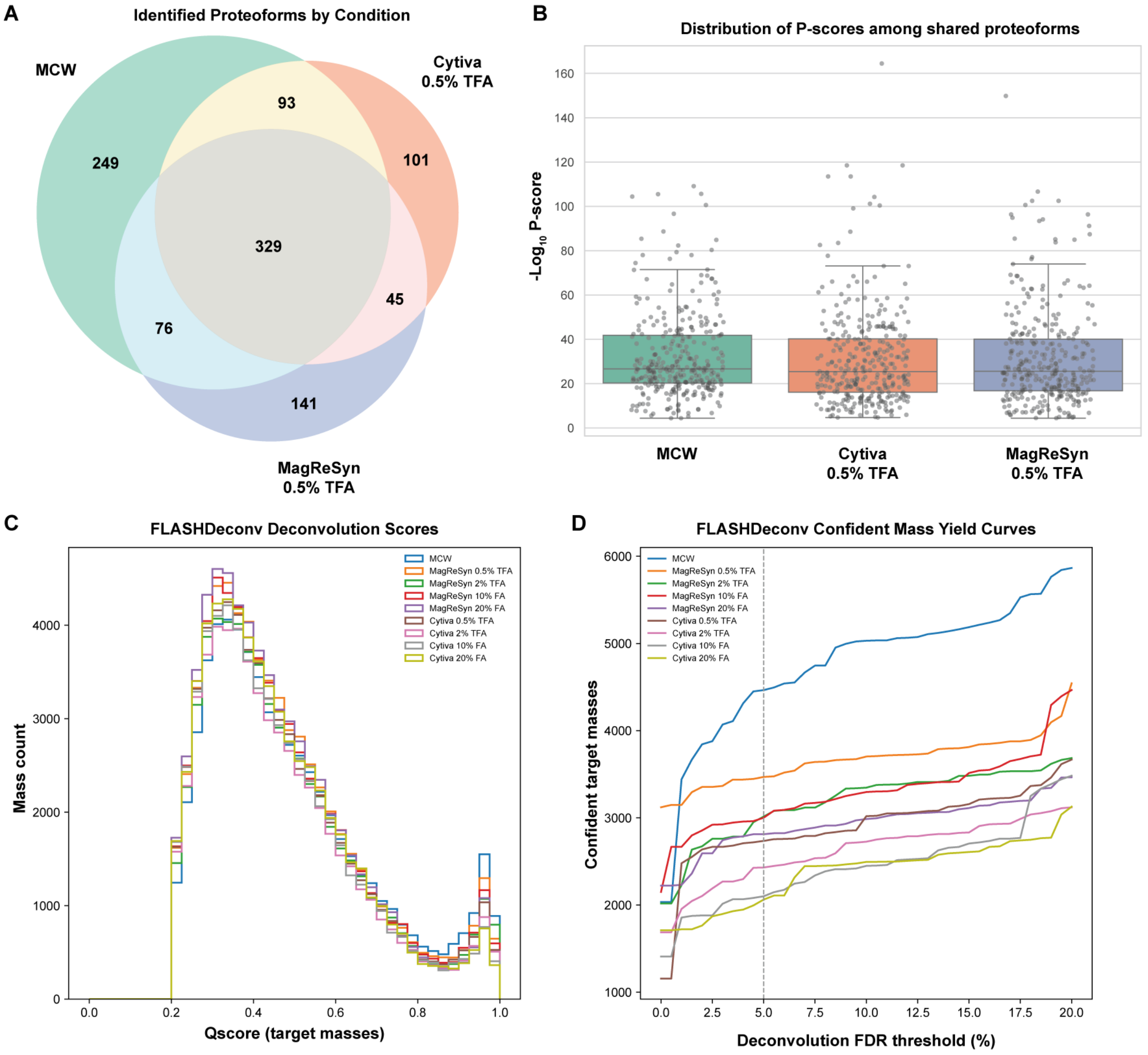


Figure S6. Proteoforms that (A) are unique and shared between experimental methods and (B) the P-scores distributions of shared proteoforms across experimental methods. (C) Distribution of FLASHDeconv Qscores. (D) Count of masses passing different FDR thresholds.

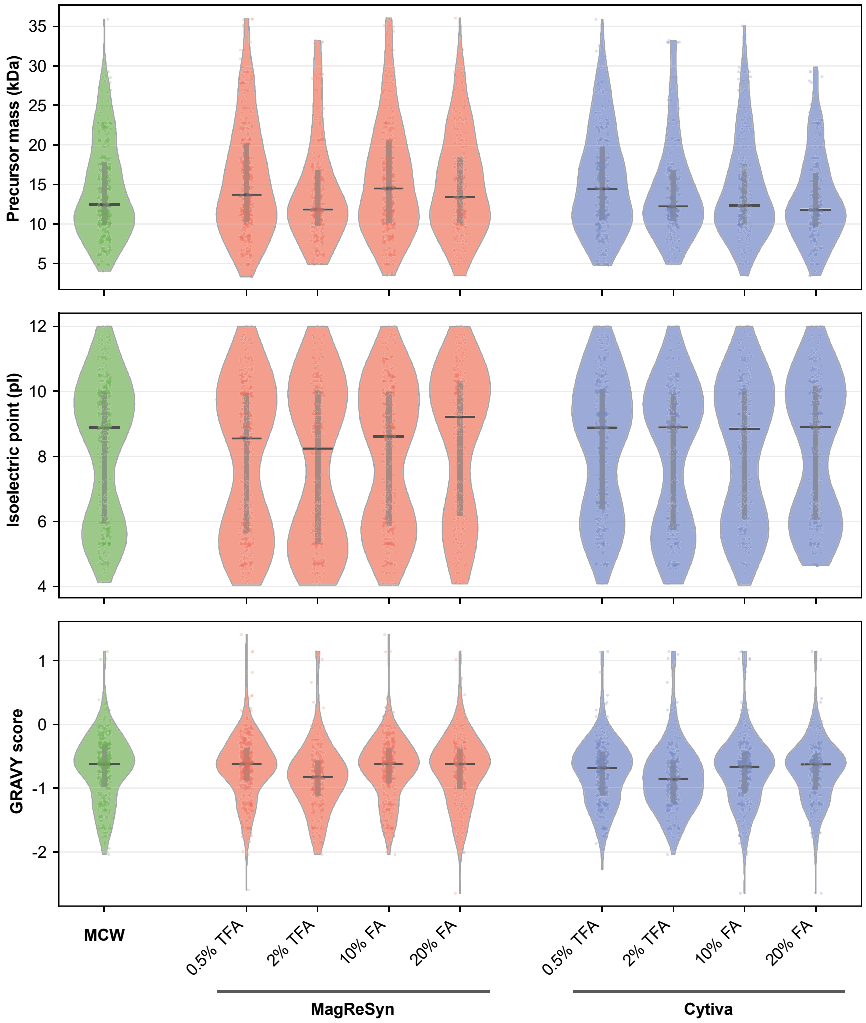


Figure S7. (A) Mass, (B) isoelectric point, and (C) GRAVY score distributions for proteoforms identified across cleanup methods, excluding histone proteins.

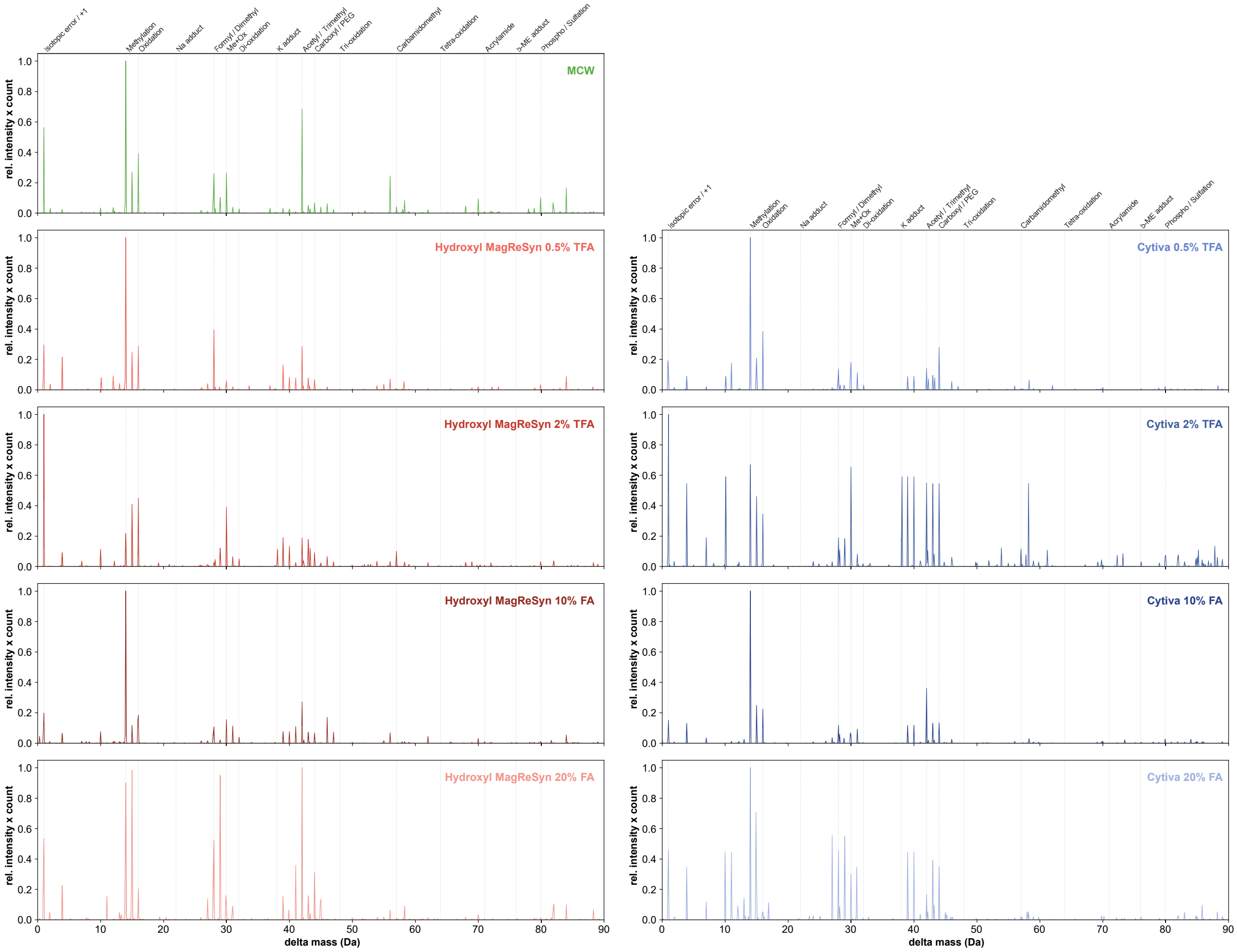


Figure S8. MSTopDiff identification of common mass shifts observed in HeLa lysate depending on sample preparation method.
